# The G1/S transition in mammalian stem cells *in vivo* is autonomously regulated by cell size

**DOI:** 10.1101/2024.04.09.588781

**Authors:** Shicong Xie, Shuyuan Zhang, Gustavo de Medeiros, Prisca Liberali, Jan M. Skotheim

## Abstract

Cell growth and division must be coordinated to maintain a stable cell size, but how this coordination is implemented in multicellular tissues remains unclear. In unicellular eukaryotes, autonomous cell size control mechanisms couple cell growth and division with little extracellular input. However, in multicellular tissues we do not know if autonomous cell size control mechanisms operate the same way or whether cell growth and cell cycle progression are separately controlled by cell-extrinsic signals. Here, we address this question by tracking single epidermal stem cells growing in adult mice. We find that a cell-autonomous size control mechanism, dependent on the RB pathway, sets the timing of S phase entry based on the cell’s current size. Cell-extrinsic variations in the cellular microenvironment affect cell growth rates but not this autonomous coupling. Our work reassesses long-standing models of cell cycle regulation within complex metazoan tissues and identifies cell-autonomous size control as a critical mechanism regulating cell divisions *in vivo* and thereby a major contributor to stem cell heterogeneity.

## Introduction

Cell size sets the fundamental spatial scale of all cellular processes. It delimits a cell’s biosynthetic capacity, affects the rates of cell migration and death, and influences cell fate decisions.^1–5^ Although a cell’s size is generally proportional to the amounts of its biosynthetic machinery and the volumes of its organelles, many proteins scale differently with cell size, leading to small and large cells becoming biochemically different.^6–8^ Therefore, each cell type likely controls its size to be within a specific range that optimizes the proteome composition for its physiological role. When cycling cells exceed their target size range, their cellular proteomes begin to resemble those of senescent cells, and they tend to permanently exit the cell division cycle.^6,9,10^

Change in cell size, especially stem cell size, has been associated with cancer and other aging-related diseases.^1,11^ However, we do not understand the mechanisms that maintain stable cell size in animals. In unicellular free-living eukaryotes, cell growth is largely limited by nutrient availability, while dedicated cell-autonomous mechanisms sense cell size and transmit this signal to control division.^12–14^ For multicellular organisms, in order to maintain tissue architecture and sustain the proper distribution of cell types, both cell growth and division processes are thought to be controlled extrinsically by extracellular signals.^13,15^ However, it is unclear how these signals are linked within a single cell to coordinate its growth and division to maintain a constant cell size. In adult stem cells that continually grow and divide, cell size affects critical stem cell functions like niche interactions, fate-specification, and tissue regeneration capacity.^10,16–18^ Therefore, it is important to understand whether cell-autonomous size control mechanisms, similar to those operating in unicellular cells, couple cell growth and cycle progression to produce the remarkable uniformity of stem cell sizes we see *in vivo* (**Figure 1A**).^19^

**Figure 1.**
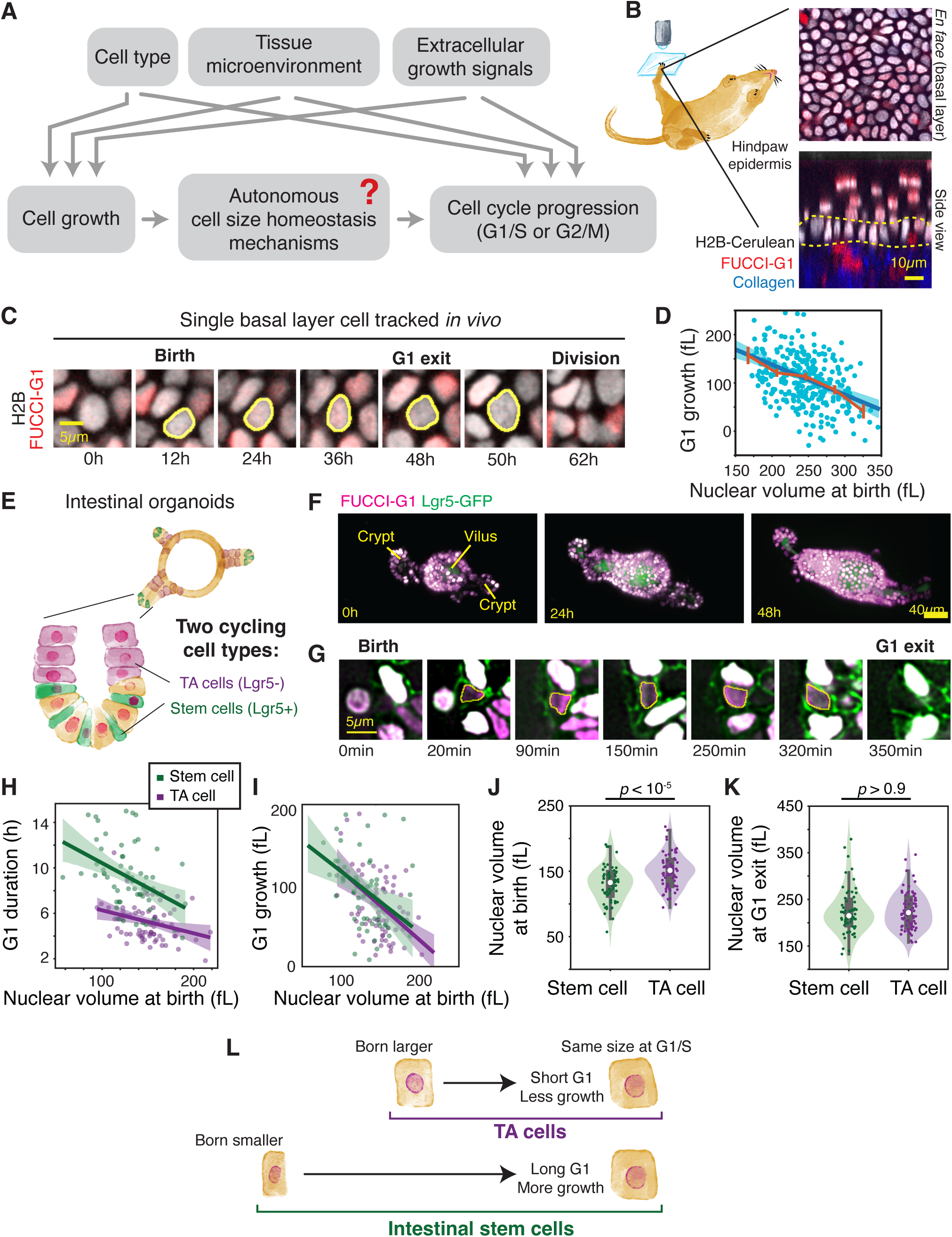
Cell size-dependence of the G1/S transition is conserved across mammalian epithelial cells. A. Schematic of the current model of multicellular cell cycle control. How cell autonomous factors and non-cell autonomous factors are integrated to regulate cell cycle progression is not well-understood. B. Intravital imaging can directly visualize cells within the mouse hind paw epidermis. *En face* and side views are shown of an epidermis expressing the *K14-H2B-Cerulean* nuclear marker and the *FUCCI-G1-mCherry* cell cycle phase reporter. Dotted yellow line indicates the location of the basal layer, where stem cells reside. C. An example of a single basal layer stem cell tracked from birth to division. D. Epidermal stem cell birth size is inversely proportional to the amount of cell growth experienced during G1 phase. Binned medians ± SEM are shown in red. Slope = −0.63. N = 2 mouse, 254 cells. E. Schematic of the tissue architecture of mouse intestinal organoids. Lgr5+ intestinal stem cells reside at the bases of intestinal crypts, while Lgr5-TA cells are located towards the neck regions. F. Maximum intensity projection images of an intestinal organoid expressing *Lgr5-GFP* and *Cdt1-mKO2* (FUCCI-G1) imaged every 10 minutes for over 36 hours using light-sheet microscopy. G. An example of a single Lgr5+ intestinal stem cell tracked from birth to G1 exit. Segmentation of nuclear shape is shown in yellow. A characteristic z-slice is shown. H. The nuclear volume at birth of intestinal stem cells and TA cells is inversely correlated with G1 duration. Pearson’s *R* = −0.42 for stem cells and −0.31 for TAS cells. N = 3 organoids, 71 stem cells, 59 TA cells. I. The nuclear volume at birth of intestinal stem cells and TA cells is inversely correlated with the amount of nuclear growth during G1 phase. Slope = −0.75 for stem cells and −0.85 for TA cells. J-K. Nuclear size at birth (J) and at G1 exit (K) is quantified for intestinal stem cells and TA cells. Median is shown in white. Shaded area is a violin plot. L. Schematic of cell size control. Cells that are born smaller (e.g., intestinal stem cell) will spend longer in G1 phase so they can grow proportionately more compared to larger-born cells (e.g. TA cell). As a result, smaller- and larger-born cells will progress through the G1/S transition at similar cell sizes. Linear regression and 95% confidence intervals are shown as a solid line and shaded area, respectively. *p*-values are from two-sample T-tests.

Previous work identified a class of cell-autonomous size control mechanisms in which the concentration of a cell cycle inhibitor becomes diluted by cell growth to preferentially trigger cell cycle progression in larger cells.^20–23^ These diluted inhibitors include the retinoblastoma protein RB1 in human cells, which binds and inhibits the E2F cell cycle transcription factors to inhibit S phase entry.^21,24^ Although inhibitor-dilution was identified as a cell size control mechanism in some cultured human cells, its functional relevance – and the general importance of size regulation at the G1/S transition – has recently been called into question. This is because a large panel of cell lines cultured *in vitro* do not strongly control their size at the G1/S transition.^25–28^ Moreover, cells *in vitro* are 3-4 fold larger than their *in vivo* counterparts and cycle much faster.^29,30^ The lack of physiological and phenomenological similarity between metazoan cells *in vitro* and *in vivo* has precluded our ability to use *in vitro* models to dissect what mechanisms coordinate cell growth and cell cycle progression *in vivo*.

Here, we use intravital time lapse imaging to examine cell size control in the adult mouse epidermis, a highly proliferative tissue where both cell and tissue size are maintained. We focus on adult stem cells because the cell-extrinsic signals in homeostatic tissues are likely less complex than those in dynamically growing embryonic tissues. Furthermore, methods now exist to directly image stem cells in the mouse epidermis *in vivo*. We previously showed that for epidermal stem cells, unlike *in vitro* cultures, the rate of G1/S transition is very sensitive to cell size.^31^ In this work, we tracked the growth and division of single epidermal stem cells alongside changes in the cellular microenvironment to find that cell size is the major determinant of the G1/S transition. Cell-extrinsic factors influence single cell growth rates, but play a lesser role in directly determining G1/S timing. The RB pathway is crucial for cell-autonomous size control *in vivo*, which accounts for most of the heterogeneity in stem cell cycle timing. Taken together, our work builds a model where cell growth rate is influenced by non-cell autonomous factors, while a cell-autonomous mechanism, dependent on the RB pathway, sets the timing of S phase entry based on current cell size.

## Results

### Cell size control occurs at the G1/S transition across multiple mammalian epithelial stem and progenitor cell types

To test size control models *in vivo*, we need to measure and track the size of growing cells in a living animal. The mouse epidermis is ideal for these purposes. The interfollicular epidermis is made up of multiple layers of epithelial cells, with stem cells residing at the basal-most layer. Epidermal cells are close enough to the surface so that two-photon imaging can be used to directly visualize growing and dividing cells, and the same cells can be imaged over multiple days in living animals (**Fig 1B**).^32,33^ We used a *K14-H2B-Cerulean* marker to measure nuclear size, and inferred cell cycle phase using the *FUCCI-Cdt1-mCherry* (FUCCI-G1) cell cycle reporter.^32,34^ A novel image analysis pipeline was then developed to segment 3D nuclear morphology and track individual cells from their birth to division (**Fig 1C; Supplemental Figure S1**). We examined growing and dividing epidermal stem cells from the mouse paw and found results similar to those we previously found when analyzing a similar dataset acquired by the Greco laboratory.^31,32^ Using nuclear volume as a proxy for cell size, we show that a cell’s nuclear size at birth is inversely proportional to the amount of nuclear growth the cell does during the G1 phase, indicating that cells accurately control their size at the G1/S transition (slope = −0.63; **Fig 1D**). This contrasts with the much less stringent cell size regulation reported for individual cells grown in culture (slope = −0.24 for HT29).^27^

To test the generality of the model that cell size control of mammalian epithelial cells predominantly takes place at the G1/S transition, we examined intestinal organoids. Epithelial cells in these organoids closely resemble their *in vivo* counterparts in terms of their size, architecture, and transcriptional state.^30,35^ In the intestinal epithelium, there are two main proliferative cell types, the Lgr5+ intestinal stem cells and the Lgr5-transit-amplifying (TA) cells (**Fig 1E**). We used light-sheet microscopy to image the growth of organoids expressing the stem cell marker *Lgr5-DTR-GFP* and the G1 phase marker *Cdt1-mKO2* (**Fig 1F**).^36,37^ We then tracked the growth of single stem or TA cells within the organoid bud region (**Fig 1G, Supplemental Fig. S2A,B**).^38^ Using nuclear volume as a proxy for cell size (**Supplemental Figure S2C,D**), we find that both stem cells and TA cells would wait longer to grow more in the G1 phase if they were born smaller (**Fig 1H,I**). In both cell types, the relationship between birth size and the amount of G1 growth lies on nearly the same line, suggesting that the cell size threshold needed to trigger the G1/S transition is the same in both cell types. Indeed, we found that despite differences in average growth rates (**Supplemental Fig S2E**) and average birth size across stem and TA cells (**Fig 1J**), both cell types enter S phase at the same average size (**Fig 1K**). Taken together, these data suggest that the rate of the G1/S transition is tightly coupled to cell size across multiple epithelial cell types, and that within the intestinal lineage, the cell size threshold required for the G1/S transition remains the same even when stem cells start to differentiate (**Fig 1L**). Therefore, it is likely that the molecular mechanism that encodes this size threshold is not remodeled by TA cell differentiation.

### Epithelial stem cell size controlled by the RB pathway

Having confirmed that the G1/S transition is crucial for controlling cell size in multiple epithelial cell types, we tested whether the coupling between cell size and the G1/S transition rate is mediated by the RB pathway, as previously reported in cell culture work.^21^ In cultured cells, Rb1 is inactivated via phosphorylation by CDKs and by dilution (**Fig 2A**). Larger G1 cells have lower concentrations of Rb1 and are therefore more likely to enter S phase. To test if the RB pathway also regulates cell size in an animal, we performed longitudinal intravital imaging and tracked single cell growth in animals where *Rb1*-deletion is inducible by tamoxifen or 4OH-tamoxifen (4OHT) treatment (**Fig 2B; Supplemental Figure S3A**).^39^ When we deleted *Rb1* by itself, we found minimal effects on cell size control (**Supplemental Fig. S3D-G**). However, its paralog *Rbl1* (p107) is known to compensate for *Rb1*-loss in both MEFs and the adult skin.^40,41^ When *Rb1* was deleted in a *Rbl1*−/− animal (DKO), the G1 duration no longer depended on cell birth size (**Fig 2C,D; Supplemental Fig. S4A**).^42^ In ear tissues where only *Rbl1* was mutated (SKO), basal layer stem cells that are born smaller had longer G1 durations compared to larger-born cells (**Figure 2E,F; Supplemental Movie S1**), similar to wild-type paw cells (**Supplemental Figure S3F**). However, in DKO tissues, cells have G1 phases of similar durations regardless of birth size, indicating a loss of cell size control at the G1/S transition (**Fig 2G,H; Supplemental Movie S2**). Consistent with the RB pathway being important for maintaining cell size homeostasis, the variation in cell size increased in DKO compared to SKO tissues (**Fig 2I**). In DKO tissues, mitotic figures were visible in the supra-basal layer, and the ear epidermis grew considerably thicker, although the K10 differentiation marker was not significantly different compared to SKO tissues (**Supplemental Figure S4B-D**). By day 4.5 post-4OHT treatment, when most DKO cells have finished their first division and initiated their second cell cycle (**Fig 2J**), cells started to die throughout the tissue in numbers significantly above those found in SKO tissues (**Fig 2K; Supplemental Figure S4E**). When we harvested skin tissues to stain for DNA damage markers, we found that DKO cells had significantly more 53BP1 foci compared to SKO cells, although it remains unclear whether this is caused by the uncoupling of S phase entry from cell size, or other functions of the RB pathway (**Fig 2L; Supplemental Figure S4F**). Therefore, mammalian cell size homeostasis depends on the RB pathway and may be crucial for successful genome replication.

**Figure 2.**
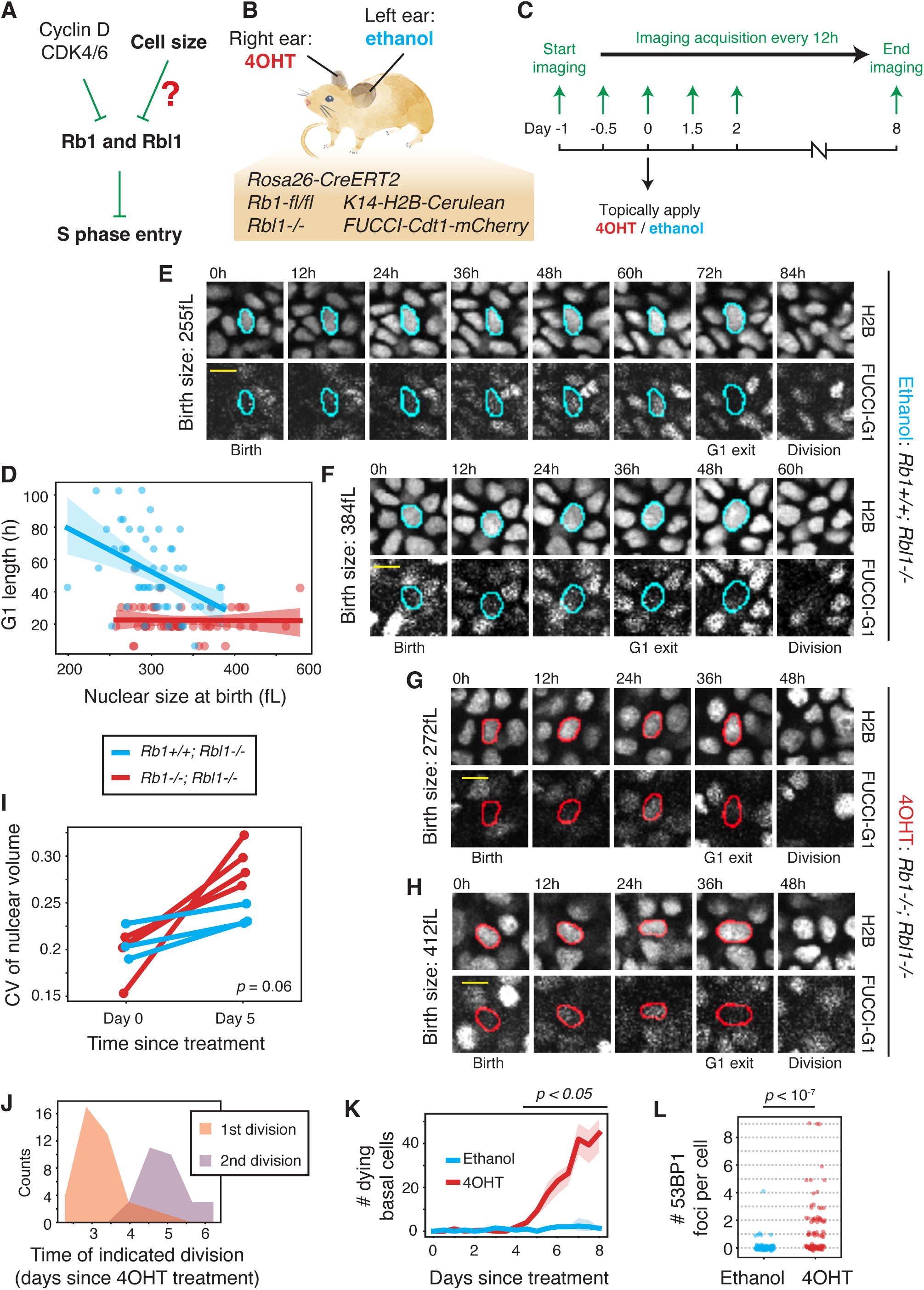
Cell size control *in vivo* depends on the RB pathway. A. Simplified schematic of the RB pathway. At the G1/S transition, the S phase-inhibiting activities of Rb1 and its paralog Rbl1 are repressed through phosphorylation by Cyclin D-CDK4/6 complexes as well as through dilution by cell size. B. 4OHT or vehicle was applied topically to the left and right ears of a mouse homozygous for *Rb1-floxP* and *Rbl1−/−* alleles. The mouse also bears a tamoxifen-inducible CreER and nuclear and cell cycle reporters. C. Imaging was started 24h prior to drug application and continued every 12h for 8 more days. The tissue was collected after imaging. D. The relationship between nuclear size at birth and the duration of G1 phase is shown for SKO and DKO cells. Linear regression and 95% confidence intervals are shown as a solid line and shaded area, respectively. SKO: Pearson’s *R* = −0.46, 57 cells; DKO: Pearson’s *R* = −0.01, 49 DKO cells. E-F. Representative montages of a smaller-born (E) and a larger-born (F) SKO cell. The smaller-born cell has a long G1 phase (72h), while the larger-born cell has a short G1 phase (36h). G-H. Representative montages of a smaller-born (G) and a larger-born (H) DKO cells. Both cells have G1 phases of similar duration (36h). I. The CV of nuclear size is shown for cells prior to the indicated treatment and 5 days after ethanol or 4OHT treatment. *p*-value is a T-test of ΔCV between genotypes. J. The distribution of timing is shown for when DKO cells finish their first and second rounds of cell division, relative to the initial 4OHT application. K. The number of visible dying nuclear bodies are shown for SKO and DKO tissues relative to the timing of drug treatment. Solid line is mean. Shaded region is std. N = 3 regions per genotype. L. The number of 53BP1 foci per nuclei is quantified for SKO and DKO tissues. SKO: 112 cells, DKO: 115 cells Scalebar is 10µm. *p*-values are from two-sample T-tests.

### Cell size is the dominant feature determining passage through the G1/S transition

Having established that tight coupling between the G1/S transition and cell size depends on the RB pathway, we sought to determine whether the G1/S transition is also regulated by the cellular microenvironment. For example, in the skin, it is thought that the local loss of cells in the basal layer upregulates the division of neighboring cells.^32,43^ However, it is unclear whether microenvironment variations simply modulate stem cell growth rates so they can reach the cell size threshold required for the G1/S transition faster, or whether variation in tissue signals or microenvironment state can directly exert influence on the timing of the G1/S transition itself (**Fig 3A**).

**Figure 3.**
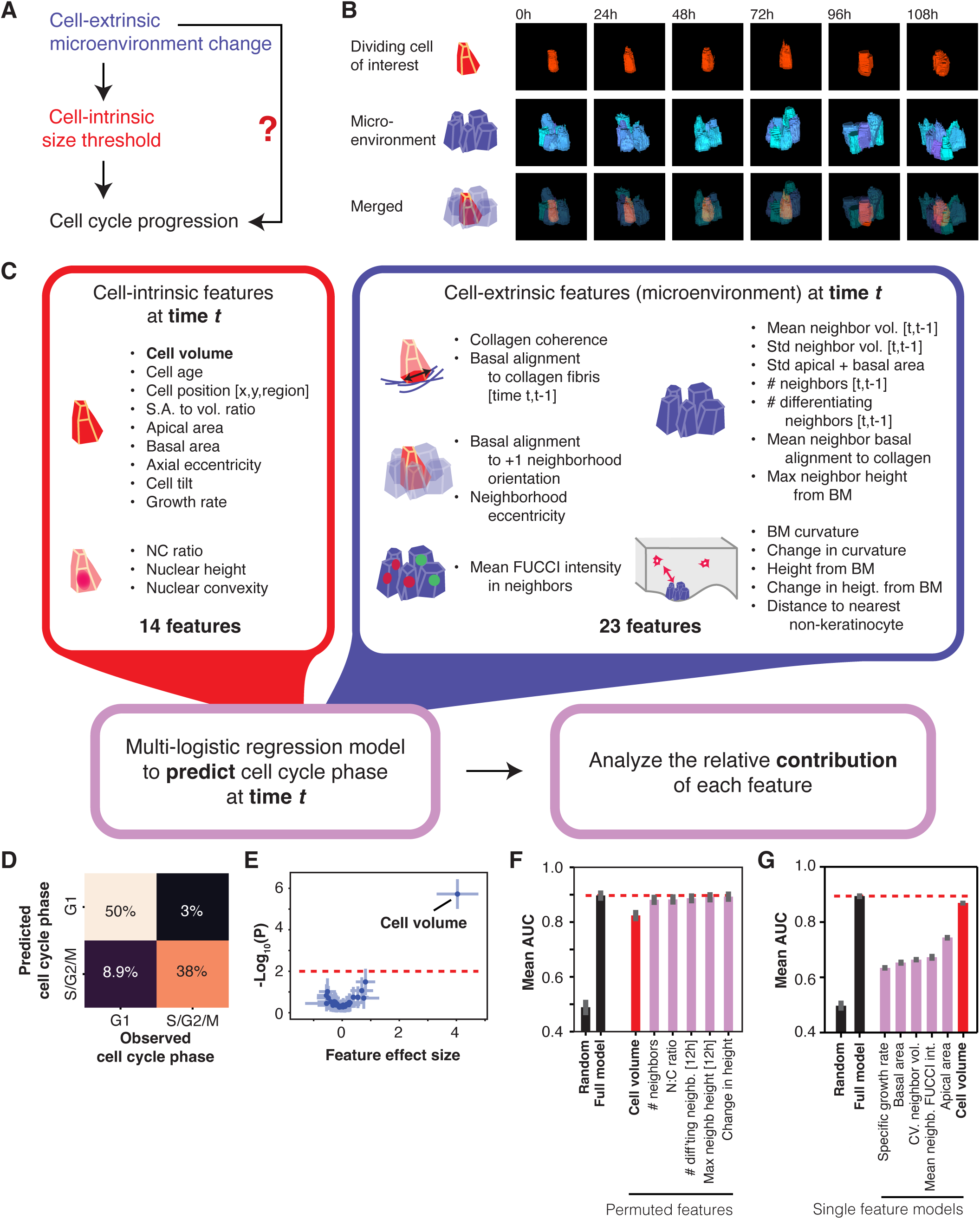
Cell size is the dominant predictor for when cells transition from G1 to S phase. A. Tissue signals or microenvironment changes could influence cell cycle progression primarily by changing the cell growth rate, making cells reach the size threshold needed for G1/S progression faster, or by directly influencing cell cycle progression without influencing cell growth. B. Example montage of the 3D quantification of cell-intrinsic geometry and cell-extrinsic microenvironment. Top row shows a single basal layer stem cell from birth to division. The middle row shows the cells that neighbor the cell of interest (microenvironment). The bottom row shows the cell of interest and its microenvironment merged. C. Predictive modeling of cell cycle variation in the basal layer tissue. Left shows a table of all the morphometric features derived from cell-intrinsic (14 features) and cell-extrinsic (23 features) geometries. These features were collated and used to train a multiple-logistic regression model to predict whether a cell will be in G1 or S/G2 phases of the cell cycle. Then, the relative contribution of each feature to the model’s statistical power was analyzed. D. Error matrix of the model showing the observed and predicted cell cycle phase classifications. The average model performance is shown for 1000 iterations with 10% randomly withheld data. N = 2 regions, 140 S phase cells, and 150 G1 phase cells subsampled randomly from 567 total G1 data points. E. The coefficient size of each feature is shown against its significance in the model, as determined by F-test for nested models where the feature was omitted from the model. Only features above significance values of α = 0.01 (dotted line) are labeled. Error bars indicate std of 1000 subsample iterations. F. The mean area under the curve (AUC) is shown for cross-validation models with the entire feature-set, completely random features, or feature-sets where one of the indicated features was randomly permuted. Only cell volume shows significant decrease in model performance when permuted. Error bars indicate 95% confidence interval from 1000 cross-validations. G. The mean AUC is shown for cross-validation models with the entire feature-set, completely random features, or models with only a single indicated feature. The cell volume-only model performs nearly as well as the full model.

To disentangle the effects of cell size and microenvironment features on the G1/S transition, we expanded our analysis of wild-type hind-paw epidermal basal layer cells growing during normal homeostasis.^31,32^ We extended our image analysis pipeline to exhaustively annotate all basal layer stem cells within the visible tissue, so that we can conduct a morphometric analysis of all cells and cell-cell contacts with high fidelity (**Supplemental Figure S5; Supplemental Movie S3**). Our methods enabled the simultaneous quantification of cell-intrinsic shape alongside quantification of the shape of that cell’s microenvironment, consisting of its immediate neighboring cells (**Fig 3B**). We measured many features of cell-intrinsic shape as well as geometries pertaining to the microenvironment, including neighborhood density, average neighbor size, and tissue curvature (**Fig 3C; Supplemental Fig. S5-6**). We used these features to build multiple-logistic regression models that can accurately predict whether a cell is in G1 or S/G2 phases of the cell cycle (**Fig 3C,D; Supplemental Fig. S7**). Surprisingly, when we analyzed the relative contribution of each feature to the model’s accuracy, we found that cell volume was the sole significant factor (**Fig 3E**). Consistent with this, we found that the model suffered in its performance only when we randomly permuted cell volume, compared to any other single feature (**Fig 3F**). Lastly, models trained on cell volume alone perform nearly as well as models trained on the whole feature-set (**Fig 3G**). Similar results were obtained when we trained the model with PCA-diagonalized feature-sets or used random forest classifiers (**Supplemental Fig. S8C-I**). Thus, reaching a cell size threshold is the dominant determinant of when a skin stem cell will enter S phase.

### The cellular microenvironment regulates growth, but not cell-autonomous size control

So far, our work supports a model in which variations in cellular microenvironment and tissue-level signals are upstream of a cell-autonomous size control mechanism. To directly test this model, we experimentally perturbed the microenvironment of epidermal stem cells and asked whether the size at which these stem cells entered S phase remained the same. We used a two-photon laser to selectively deliver a high dose of light to a single cell, and quantified the volumetric growth rates and S phase entry sizes of cells that were either within 20µm of the ablation site (neighboring) or further away (non-neighboring) (**Fig 4A; Supplemental Figure S9A-C**). Within 8-12h, the ablated cell disappeared from the tissue while the neighboring cells remained intact and growing (**Fig 4B**). When we examined the exponential growth rates derived from the growth of either neighboring or non-neighboring cells, we saw that cells that neighbored the ablation site grew significantly faster for 3 out the 4 animals we analyzed (**Fig 4C; Supplemental Figure 9D,E**). However, for all animals, the size at which cells entered S phase remained indistinguishable between neighboring and non-neighboring cells (**Fig 4D**). Therefore, the coupling of cell size to G1/S transition rates remains similar despite ablation-neighboring cells experiencing faster growth rates. These data collectively support the model in which variation in tissue signals or microenvironment states mainly modulate stem cell growth rates, but the G1/S transition decision is predominantly determined by the cell’s current size (**Fig 4E**).

**Figure 4.**
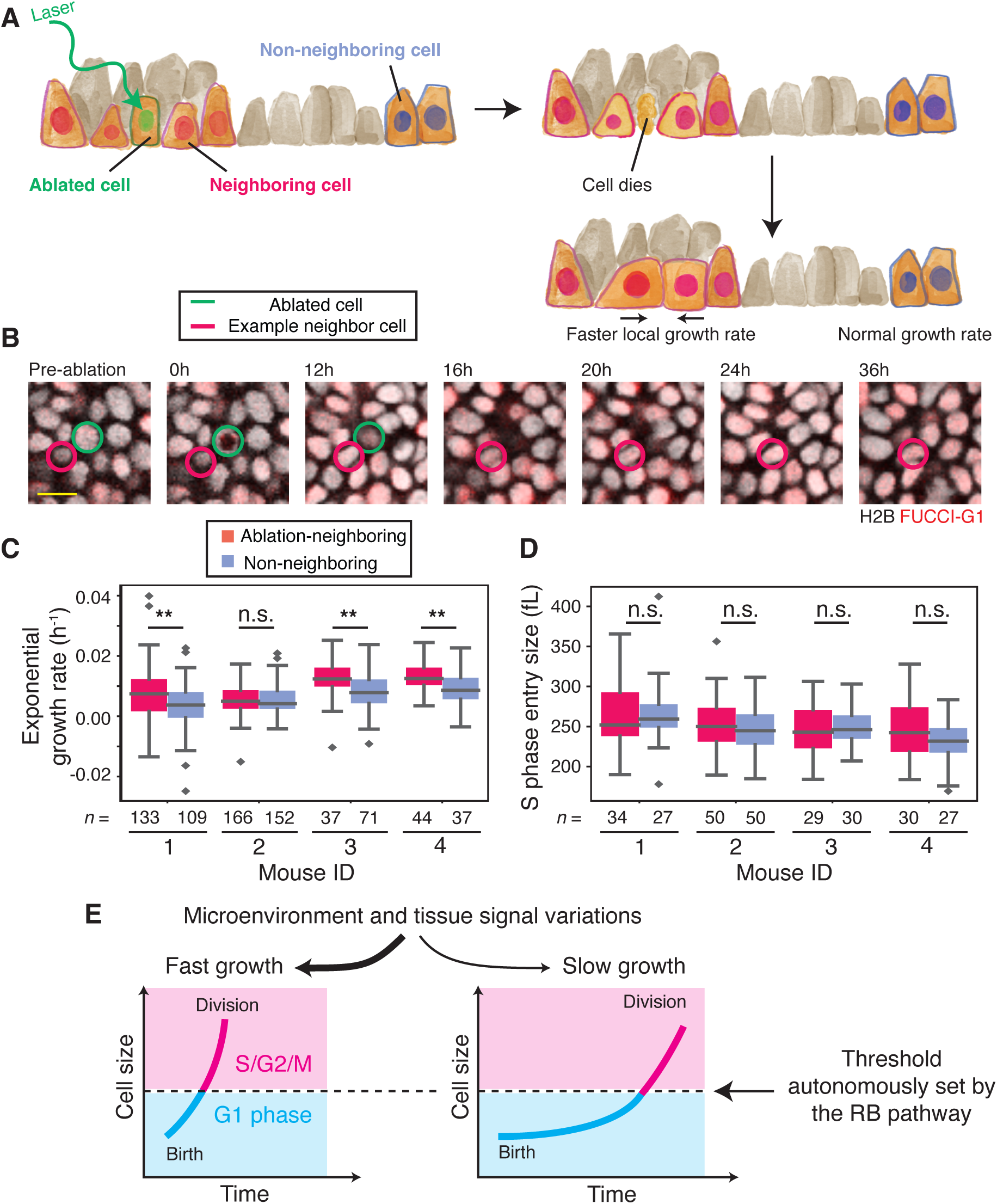
Microenvironment perturbations affect neighbor cell growth, but not cell autonomous size control. A. Schematic of cell ablation experiments to perturb the skin stem cell microenvironment. A high dose of light is delivered to a single cell nucleus, leading to cell death. The nearby neighboring cells experience faster cell growth rates compared to non-neighboring cells, which are further away from the ablation site. B. Example montage of a cell ablation time-series, showing a central ablated cell (green) and a single neighbor cell (red) tracked through time. Only a single tracked neighbor cell is highlighted for clarity. Scale bar is 10µm. C. Exponential growth rate fitted from the nuclear growth rate of ablation-neighboring and non-neighboring cells are shown for 4 mice. The midline shows the median and the box delimits the quartiles. **: *p* < 0.01, n.s.: *p* > 0.05, from two-sample T-tests. D. The nuclear size at which cells enter S phase is shown for ablation-neighboring and non-neighboring cells. n.s.: *p* > 0.2, from two-sample T-tests. E. In adult tissues, variations in tissue and microenvironment state influence the rates of stem cell growth but do not directly influence G1/S transition rates. Instead, a cell size control mechanism, dependent on the RB pathway, autonomously couples the instantaneous cell size to the G1/S transition.

## Discussion

The complexity of cell-extrinsic control in multicellular tissues has raised the question of whether any cell-autonomous size control mechanisms identified from unicellular eukaryotes and *in vitro* cell lines can couple cell size to G1/S transition rates *in vivo*. Here, we show that such cell-autonomous mechanisms are of primary importance for cell size and cell cycle regulation in multiple epithelial stem cell types, both *in vivo* and in organoids. Given that recent work also found size coupled G1/S transitions in mature MDCK monolayer cultures, cell-autonomous size control is likely a general feature of epithelial tissues.^30^

We were able to reach the surprising conclusion that cell-autonomous size control is dominant over non-cell autonomous signaling in controlling the G1/S transition through our exhaustive annotation of basal layer cells and their cell-cell contacts. By quantitatively characterizing the microenvironment over time, we could eliminate fluctuations in microenvironment geometry as proximal causes of the G1/S transition. Cell-ablation experiments further supported our conclusions that the microenvironment plays an important role in regulating cell growth but not in directly regulating cell cycle transitions. Loss of a neighbor cell upregulates stem cell growth and makes cells reach the critical size needed to traverse the G1/S transition faster, but does not change the critical size itself. Instead, the decision to enter S phase is cell-autonomously determined by a molecular coupling of the G1/S transition to cell size, likely via the dilution of RB (**Figure 4E**).^21,24^

This cell-autonomous model of how the G1/S transition is controlled *in vivo* calls into question our previous understanding of cell cycle control in the context of tissue biology. Historically, studies have inferred from fixed cells or tissues that animal cells do not exhibit cell-autonomous size control, with cell cycle progression seemingly unaffected by cell size.^44^ In these models, often based on embryonic or developmental contexts, cell cycle progression and cell growth are independently controlled by cell-extrinsic signals.^45,46^ The difference between our conclusions and previous developmental models of cell cycle control may be that during development both cell and tissue size are not kept in homeostasis, whereas in the adult skin they are. Thus, cell-autonomous size control could be a crucial mechanism that supports the maintenance of adult tissues.

Finally, we suspect that keeping efficient stem cell size homeostasis is important for maintaining tissue health throughout the organism’s lifespan, since recent work has linked stem cell enlargement to a loss of regeneration potential in the blood and intestine.^10^ Indeed, blood and skin stem cells become increasingly large with age, which is coincident with their progressively declining function.^10,47^ Therefore, understanding how cell-autonomous size control mechanisms break down during the aging process could yield new strategies for treating diseases associated with aging.

## Acknowledgments

We thank Valentina Greco, Julien Sage, and Fred de Sauvage for generously sharing their reagents, and David Gonzalez, Katie Cockburn, and Gordon Wang for their advice on experimental design. We thank Katie Cockburn, Alex Lessenger, Roel Nusse, and Lucy O’Brien for comments on the manuscript. This work was supported by NIH grants R01AR079860 to J.M.S. and K99GM138712 to S.X., a Chan Zuckerberg Initiative award to J.M.S., and a Travelling Fellowship from The Company of Biologists to S.X.. The Stanford Cell Sciences Imaging Facility provided training and usage support with RRID:SCR_017787. The Stanford Neuroscience Microscopy Service also provided training and usage support. This work used supercomputing resources provided by the Stanford Genetics Bioinformatics Service Center, supported by NIH Instrumentation Grant S10 OD023452.

## Attributions

S.X. and J.M.S. conceived and designed the experiments. S.X. generated organoid lines, acquired light-sheet and intravital imaging data, maintained mouse lines and performed crosses, designed the analysis software, and performed data analysis. S.Z. designed and performed mouse crosses and aided with mouse experimental design. G.M. and L.P. aided with the setup, acquisition, and analysis of light-sheet images. S.X. generated the illustrations. S.X. and J.M.S. wrote the manuscript.

## METHODS

### Mouse lines

All mice were handled in accordance with the guidelines of the Institutional Animal Care and Use Committee at Stanford University. The *K14-H2B-Cerulean* and *R26p-FUCCI2* mice were kindly shared by Dr. Valentina Greco at Yale University. The *Rb1-flox*, *Rbl1−/−*, and *Rosa26-CreERT2* mice were kindly shared by Dr. Julien Sage at Stanford University. The *Lgr5-DTR-GFP* mice were kindly shared by Dr. Fred de Sauvage at Genentech. Unless otherwise noted, mice used for intravital skin imaging in the study were of the genotype *K14-H2B-Cerulean*; *R26p-FUCCI2*; *Rosa26-CreERT2*; *Rb1-fl/fl*. Wild-type mice refer to mice of this genotype that were never exposed to tamoxifen. For visualizing cell cycle phases, only the *Cdt1-mCherry* portion of the *R26p-FUCCI2* reporter system was visible in our studies because the strong signal from the *K14-H2B-Cerulean* reporter masks the signal of the *Geminin-mVenus* reporter in the yellow channel. All mice imaged in the study were 3-7 months old and never used for breeding.

Table of mouse lines used in this study:

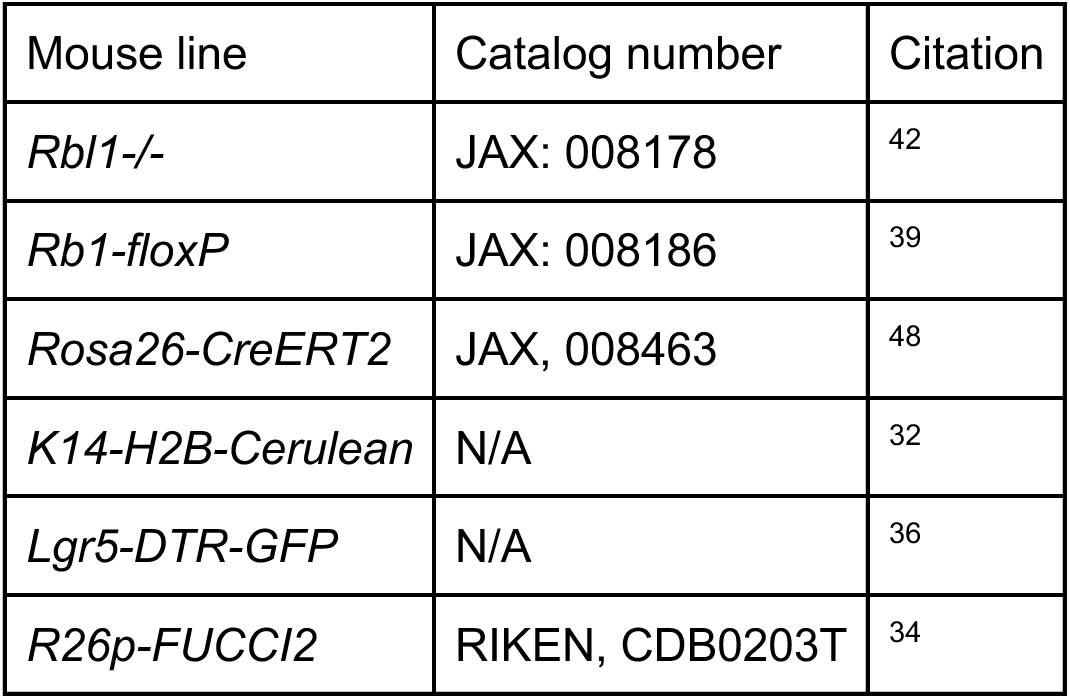

### Induction of CreER activity using tamoxifen or 4OHT

To induce CreER activity in the paw skin, 10 mg/mL tamoxifen dissolved in corn oil was introduced via intraperitoneal injection to the mouse abdomen for 3-5 days consecutively. Subsequently, the mice were imaged 4-5 weeks post-injection for *Rb1−/−* single mutant experiments.

For DKO experiments, topical application of 4OHT to the thinner ear skin was instead used to directly compare SKO and DKO tissues in the same animal. A slurry of 4OHT was applied topically to the ear skin, consisting of a solution of ethanol and 2mg/mL of 4OHT mixed with 1g of petroleum jelly (Vaseline). 100mg of the 4OHT slurry was then applied on top of the ear of an isoflurane-anesthetized mouse, let incubate for 30 min, and washed thoroughly. As a control, ethanol-only slurry was applied to the other ear. The mouse was imaged before and after the 4OHT treatment as described in the main text (**Figure 2C**).

### Longitudinal intravital imaging

The longitudinal imaging of the same mouse skin region was performed based on protocols previously described.^32,33,49^ At least one week prior to imaging experiments, the mouse ear was depilated using a depilating cream (Nair). During imaging, the mouse was kept under anesthesia by isoflurane inhalation through a nose cone and kept warm by a rodent heating pad. The mouse paw or ear skin was immobilized between a custom-made stage arm and another stage arm holding a coverslip. The coverslip pressure was kept to a minimum to immobilize the tissue without deforming it. To find the same tissue regions over time, extensive cartographic notes were taken noting the idiosyncratic features of the skin itself, including folds, hair follicle patterns, and blood vessel patterns. The exact matching tissue region was confirmed by close comparison between the underlying dermal collagen signal from session to session. Images were taken on a Prairie View Ultima IV (Brucker) scope equipped with a DeepSee II tunable laser (Coherent). The sample was imaged with either an Olympus XLUMPLFLN 20x water-dipping objective (NA = 0.95) or an Olympus LMUPlanFl/IR 40x water-dipping objective (NA = 0.8). For visualizing Cerulean, the excitation laser was tuned to 920 nm or 940 nm and both 460/50nm and 525/50nm bandpass filters were used. To visualize mCherry, the excitation laser was tuned to 1020nm or 1040nm and a bandpass filter of 595/50nm was used. Collagen fibrils were visualized with 460/50nm bandpass filter using 920-940nm excitation or 525/50nm bandpass filter using 1020-1040nm excitation.

### Segmenting and tracking single cells in longitudinal skin images

A custom image analysis pipeline was built to register, segment, and track epidermal stem cells in the basal layer, in part based on previous work.^31^ The longitudinally acquired images were first registered using the second-harmonic generation collagen signal, since collagen fibrils rarely move during skin homeostasis. Cross-correlation was used to identify the respective z-positions between two time points that corresponded to each other, and a Python wrapper of StackReg was used to register the two tissue volumes.^50^ If bending or warping of the tissue impeded successful tracking of cells, then, BigWarp in FIJI was optionally used to nonrigidly warp the tissues using collagen landmarks.^51,52^ Nuclear image volumes were first treated with local contrast equalization using equalize_adapthist from scikit-image. Cellpose was then used in 3D stitching mode to segment the nuclear volumes in 3D, using the pretrained nuc model.^53^ Independently, single cells were manually tracked from birth to division (or only until G1 exit) using MaMuT and BigDataViewer in FIJI.^54,55^ Then, the coordinates of the single cell tracks were mapped onto the registered movie, and the corresponding nuclear segmentations were collated. To preserve cell volume, if BigWarp was used to register the movie, the nonrigid transformations were reversed and tracking coordinates mapped back onto untransformed images. napari was used for manually inspecting, curating, and editing the 3D segmentations.^56^ Cell cycle timings were manually annotated. A final script was used to collate the segmented and tracked cells with the cell cycle timing annotations to generate time-series measurements for every tracked epidermal stem cell. See **Supplemental Figure S1** for a graphical representation of the pipeline.

Throughout the paper, except for in Figure 3, nuclear volume was used as an approximate proxy for cell volume, since we have previously reported that nuclear and cell volumes are linearly proportional to each other.^31^ For DKO cells, because the duration of G1 phase was rendered very short (median = 24h) by the mutations, we were unable to accurately estimate the precise size at the G1/S transition with our 12h sampling rate. However, the change in the duration of G1 duration compared to SKO cells at different sizes is well-estimated by our sampling.

### Lentiviral generation

The FUCCI-Cdt1-mKO2 reporter construct was cloned into the CSII-EF-MCS lentiviral vector backbone under a constitutive EF1α promoter.^57^ The CSII vector, the lentiviral packaging vector dr8.74, and the envelope vector VSVg were transfected into HEK 293T cells by PEI (1mg/mL, Sigma-Aldrich). The 293T cells were maintained in DMEM supplemented with 10% FBS at 37C with 5% CO_2._ 48-60 hours later, 10mL of the lentivirus-containing medium was collected and filtered through a 0.45µm filter. The viral supernatant was concentrated by centrifugation at 50,000 *g* for 2:20 hours at room temperature. The viral pellet was dried and resuspended in 0.5mL IntestiCult medium (StemCell Technologies) supplemented with 10µM Y-27632 and 2.5µM CHIR-99021.

### Intestinal organoid culture generation, maintenance, and engineering

Intestinal organoids were generated from a 3 month old male mouse bearing the Lgr5-DTR-GFP allele by methods established previously.^58,59^ Briefly, the proximal 15cm section of small intestine was collected, opened longitudinally, and cut into small ∼2 mm sections. The intestinal pieces were washed 20 times in PBS by pipetting up and down in 10mL pipettes. The pieces were gently dissociated with EDTA and resuspended with PBS and 0.1% BSA and shaken vigorously. Four fractions of supernatant were collected, strained through a 70µm cell strainer, and each fraction visually examined for intact crypt morphology. The best crypt fractions were concentrated and resuspended in a mixture of 50:50 IntesitCult medium (StemCell Technologies) to Matrigel (Corning), and plated as dome-shaped droplets within IntestiCult medium. For the maintenance of organoid cultures, IntestiCult media was changed every 2-3 days, and the culture was split by dissolving the Matrigel using ice-cold PBS and using mechanical dissociation through a narrow glass or plastic pipette tip. All organoids were cultured at 37C with 5% CO_2_.

To express FUCCI-Cdt1-mKO2 in organoids, organoids were infected with lentivirus using previously established methods.^60^ Briefly, organoid cultures were digested into single cell suspensions by incubating in TrypLE supplemented with 10µM Y-27632 at 37C for 3-5 min or until single cell dissociation is confirmed by visualization. The cells were strained with a 40µm cell strainer and washed with DMEM/F12. Cells were then resuspended in 250µL of IntestiCult medium supplemented with 10µM Y-27632 and 2.5µM CHIR-99021 that also contains the lentiviral preparation. Then, two wells from a 24-well plate were bottom-coated with Matrigel and allowed to solidify. The cell-virus suspension was plated atop the Matrigel and incubated overnight at 37C. Next morning, the supernatant in the well containing viruses and dead cells were carefully discarded. An additional layer of Matrigel was then overlaid on top of the live cells that have attached to the bottom Matrigel layer. The whole well was then incubated with IntestiCult medium supplemented with 2.5µM CHIR-99021. The CHIR-99021 was withdrawn 2-3 days later. The organoids were allowed to grow out from the primary viral induction. Single organoids expressing the FUCCI reporter were hand-picked and re-cultured from single cell suspensions for at least five passages before experiments.

### Light-sheet microscopy and analysis

Organoid cultures used for imaging were transitioned to ENR medium for at least 3 days. Organoid cultures were replated into a 1:1 mixture of Matrigel and ENR medium containing Advanced DMEM/F12 supplemented with 1x Penn/Strep, 1x GlutaMax (ThermoFisher), 1x B27 (Life Technologies #17504044), 1x N2 (Gibco #A1370701), 1mM n-Acetylcysteine (Sigma), 50ng/mL recombinant mouse EGF (Gibco #8044), 100ng/mL recombinant mouse Noggin (R&D Systems #1967-NG), and 500ng ng/mL recombinant mouse R-Spondin1 (R&D Systems #7150-RS).

Preparation of organoids for light-sheet microscopy followed methods previously established.^38^ Organoids were mechanically dissociated by triturating through a P100 or P200 pipette tip, and resuspended in a 60:40 mixture of Matrigel and ENR medium. The mixture was placed as 5µL drops into a custom imaging chamber, and incubated with ENR medium for 1 additional day before being transferred into the microscope chamber. A LS1-Live dual illumination and inverted detection light-sheet microscope by Viventis Microscopy Sàrl was used. Organoids with visible budded regions were selected and imaged every 10 min for up to 3 days, with z-spacing of 0.67µm. The medium was exchanged every day.

For analysis, lightsheet movies were first deconvolved using Huygen Software by Scientific Volume Imaging, utilizing empirically measured point spread functions. The movies were then indexed using BigDataViewer to facilitate image viewing on small-memory computers. MaMuT was used to manually track single cells within the budded region of organoids. The birth frame, G1 exit frames, and spatial locations of single cells were then used to select cells for manual segmentation and nuclear volume quantification. For each cell, an estimate of its nuclear volume at birth and at G1 exit was averaged from the first and last 2-3 frames in which the FUCCI-G1 nuclear signal was visible above background, respectively. To quantify nuclear growth rates, nuclear volume was segmented for each cells every 30 min, and a smoothing cubic spline was used to fit the growth curves to generate estimates of instantaneous growth rates. Stem cells were identified by a clear border of Lgr5-GFP signal all around a cell’s nucleus. TA cells were identified by the absence of Lgr5-GFP and if a cell exited G1 phase, indicating active cycling.

### Immunofluorescence measurements in intestinal organoids

To measure the relationship between nuclear volume and cell volume, intestinal organoid cultures growing in ENR medium in 4-chamber LabTek chamber slides were fixed with 4% paraformaldehyde for 30 min at room temperature. They were blocked with 1% BSA and 2% Triton-X in PBS and stained overnight at 4C with primary antibodies. Then, they were washed three times in PBS and stained overnight at 4C with secondary antibodies. Primary antibodies: β-Catenin (BD Transduction Laboratories, monoclonal mouse, 1:400). Secondary antibodies: Goat anti-Mouse IgG (H+L) Highly Cross-Adsorbed Secondary Antibody, Alexa Fluor™ Plus 647 (Invitrogen #A32728, 1:1000). Lastly, samples were stained with DAPI and stored in PBS. Finally, the organoids were imaged using a Zeiss 880 confocal microscope. napari was used to manually segment nuclear and cell volumes.

### Whole mount epidermal immunofluorescence

Mouse epidermal tissues were prepared for whole mount imaging of interfollicular tissues according to established dissection methods.^49^ Mice were sacrificed by overdose of isoflurane anesthesia. The ear was depilated using Nair postmortem. The ear and paw were collected and the entire skin was dissected away from muscle and bone tissues. The skin was then incubated floating on top of a solution of PBS and 5mg/mL Dispase for 15-20 min at 37C. Superfine forceps were used to manually dissect the epidermal layer under a dissecting scope. The epidermal tissue was then fixed by floating atop a solution of PBS and 4% PFA for 30 min. The fixed tissue was then washed and blocked (5% normal goat serum, 1% BSA, 0.2% gelatin, and 2% Triton-X in PBS) before being incubated overnight with a primary antibody. Then, the samples were washed three times and incubated with a secondary antibody. Primary antibodies: phospho-RB1[S807/S811] (Cell Signaling Technology #9308, monoclonal rabbit 1:200); 53BP1 (Novus Biologicals #NB100-304, monoclonal rabbit 1:200); Keratin 10 (Biolegends #19054, polyclonal rabbit 1:200). Secondary antibodies: Goat anti-Rabbit IgG (H+L) Highly Cross-Adsorbed Secondary Antibody, Alexa Fluor™ Plus 647 (Invitrogen #A32733, 1:1000). Stained samples were washed three times in PBS and then mounted in Vectashield Antifade Mounting Medium (Vecta Laboratories #H-1000-10) on glass slides. Finally, the samples were imaged using a Zeiss 880 or Zeiss 980 confocal microscope. 53BP1 foci were manually counted using napari.

### Measurement of the epidermal stem cell microenvironment

A custom image analysis pipeline was built to densely annotate cell and nuclear shapes in order to measure the microenvironment surrounding stem cells of interest. We used existing imaging data published by the Greco group, which were derived from movies of basal layer stem cells growing in a mouse expressing *K14-H2B-Cerulean*, *FUCCI-mCherry*, and *K14-Actin-GFP*.^32^ We had previously sparsely annotated the majority of all stem cells that complete a cell cycle within the 7 day duration of the movie.^31^ For this study, we built new tools to annotate each dividing cell’s microenvironment by quantifying the rest of the tissue.

To densely annotate the rest of the cells in the tissue, we used Cellpose running in 3D stitching mode to segment the images using the pretrained ‘nuc’ and ‘cyto2’ models (**Supplemental Figure S5A,B**). To find the basal layer cells, automatically annotated the location of the basement membrane (BM). We used a Gaussian blurring filter with large sigmas (25-30 for xy, 5-10 for z) to blur the H2B-Cerulean images. The first-derivative of the profiles of the blurred intensity were generated at each x,y pixel position, and the z-slice that maximized the first-derivative magnitude was determined as the location of the BM (**Supplemental Figure S5C**). Using the BM location, basal layer cells could initially be identified. Then, napari was used to fine-tune basal cell identification, correct segmentation errors, as well as add back basal cells that were missing from the initial segmentation. For the purposes of this analysis, the *K14-ActinGFP* signal was examined manually and any cell that had a visible basal footprint immediately above the BM were counted as a ‘basal layer cell’, even if that footprint was very small and likely in the process of delamination. The cell’s height from the BM was recorded to capture the spectrum of the delamination process.

The cortical channel was then carefully examined to determine which cells shared a cell-cell interface and the information encoded in a 2D contact image (**Supplemental Figure S5D**). Finally, the 3D cortical segmentation was manually inspected and error-corrected using napari. To characterize apical and basal cell area, the original image volume was re-sliced to generate a flattened version of the tissue (**Supplemental Figure S5E**). The cell’s apical and basal areas were calculated as the average area of the 3 top-most and bottom-most slices, respectively.

Finally, since collagen orientation is thought to guide keratinocyte growth in development, we also measured the properties of the dermal collagen fibrils in relation to basal layer cells.^61^ To do so, gradient images were calculated from a flattened version of the collagen signal 3µm below the basal layer and smoothed with a Gaussian filter (sigma=4px). For each cell, its basal footprint was created from the flattened reslice of the tissue, and a local average Jacobian matrix was calculated based on the local gradients. Then, the eigenvalues and eigenvectors of the local Jacobian matrix were found. The orientation of the fibrils was defined as the directionality of the principal eigenvector, while a ‘coherence’ metric was defined as the ratio of the two eigenvalues, which reports the degree of alignment amongst the local fibers (**Supplemental Figure S5F**). A triangular mesh model of the basal layer surface was generated using the library trimesh and used to calculate the local tissue curvature (**Supplemental Figure S5C**).^62^ Finally, cell bodies that do not express the *K14-H2B-Cerulean* reporter are often visible throughout the epidermal layer based on a coherent cell-like shadow. These dark cell bodies likely correspond to resident macrophages.^63^ The locations of these cells were annotated and incorporated into the model (**Supplemental Figure S5C**).

For each basal cell nuclear and cell annotation, regionprops from scikit-image was used to generate individual object statistics, including cell and nuclear shapes and positions.^64^ In addition, for each previously tracked dividing stem cell of interest, statistics about the neighboring cells were calculated or collated in custom Python scripts. Although more than 130 features were generated, ultimately only a total of 37 features were kept in the analysis. An example cell and six of its microenvironment statistics are shown in **Supplemental Figure S6**.

Many co-correlated features were eliminated or combined into ratiometric features. For example, while cell volume was kept, cell surface area (SA) and nuclear volume were transformed into SA-to-volume and nuclear-to-cytoplasmic volume ratios. The resulting feature-set has relatively low covariance amongst the individual features and a reasonable condition number (C = 34.5, using the minimum covariance determinant as the estimator of the covariance matrix) (**Supplemental Figure 7A**). A similar condition number (35.3) was obtained using the empirical covariance matrix. When principal component analysis (PCA) was performed using PCA from scikit-learn, the variance was reasonably evenly weighted amongst the components, with the top 3 components combined only explaining <25% of total variance (**Supplemental Figure 7B**).^65^ Therefore, we decided to directly use the 37 features for constructing statistical models.

A full table of the features used in the model and a brief description is presented:

| Feature | Description, if necessary |
| --- | --- |
| Cell age |  |
| x,y,region | Cell location and regionID |
| Cell volume |  |
| Apical area | Average area of the top three z-slices of the cell segmentation |
| Basal area | Average area of the bottom three z-slices of the cell segmentation |
| SA to vol ratio | Cell surface area / cell volume |
| NC ratio | Nuclear volume / cell volume |
| Nuclear solidity | Nuclear volume / Volume of convex hull around nuclear shape |
| Cell eccentricity | The ratio of the magnitude of the largest moment of inertia to the second largest moment |
| Axial angle | The angle between the cell's largest moment of inertia to the z-axis |
| Relative nuclear height | The difference between the nuclear centroid's z-position and the cell centroid's z-position |
| Neighborhood eccentricity | The 2D eccentricity of the combined basal footprints of the cell's current neighbors, reflecting how 'round' or 'spindly' the local neighborhood is within the 2D basal layer sheet |
| Cell alignment to neighborhood orientation | The cosine of the angle between the cell's basal footprint's orientation and the orientation of the combined basal footprint of the cell's current neighbors |
| Cell alignment to neighborhood orientation [t-1] | Same as above but from the previous time point |
| Mean curvature | The mean curvature of the local neighborhood in 3D |
| Change in curvature | The difference in curvature from time t to t-1 |
| Collagen coherence | How co-aligned the dermal collagen fibrils are below the current cell's basal footprint |
| Alignment to collagen | The cosine of the angle between the cell's basal footprint orientation and the orientation of the collagen fibrils |
| Distance to nearest non-keratinocyte |  |
| Height to BM | The distance from cell centroid to the BM |
| Change in height | The difference in height (previous item) from time t to t-1 |
| Number of neighbors [t,t-1] | Number of neighbors touching the central cell [at time t and t-1] |
| Number of differentiating neighbors [t, t-1] | The number of neighbors whose centroids were higher than 3.5μm from the BM and therefore is likely in the process of differentiating [at time t and t-1] |
| Mean neighbor volume [t,t-1] |  |
| Std neighbor volume [t,t-1] |  |
| CV neighbor apical area |  |
| CV neighbor basal area |  |
| Max neighbor height to BM [t,t-1] |  |
| Mean neighbor collagen alignment [t,t-1] |  |
| Mean neighbor Fucci intensity |  |
| Specific growth rate | The cell's current growth rate normalized by its current volume |

### Statistical modeling of cell cycle phase

Statistical models were built using the Python libraries scikit-learn and statsmodels.^65,66^ For logistic regression, either logit from statsmodels or LogisticRegression from scikit-learn were used. For random forest classification, RandomForestClassifier from scikit-learn was used. Prior to training, all features were standardized to have a mean of 0 and standard deviation of 1.

For classification models aiming to predict whether cells have or have not passed the G1/S transition, we only kept data from the single movie frame immediately following the transition into S phase. This avoids late G2 and mitosis phase cells lending too much statistical power to the classifier in predicting the earlier G1/S transition. As this analysis leads to a large imbalance between G1 and S/G2 training examples, for training, the G1 class was randomly subsampled (without replacement) from a total of 567 examples down to 150 instances, which was comparable to 140 instances of S/G2 cells. This subsampling was randomly repeated each time a new model is trained. For all models, unless indicated otherwise, model metrics are reported as the mean model performance as evaluated on 10% of the data that were withheld from the training.

To analyze the contribution of each feature to the statistical power of the models, we re-evaluated the model performance (AUC) when each feature was dropped from the model, or when only a single feature was included in the model. To estimate the performance of a null model, we generated a random-feature model using a feature-set of the same size, but where each feature was instead drawn randomly from a Gaussian distribution with mean 0 and sigma 1.

To evaluate whether cell volume remains the leading predictor of the G1/S transition with other model structures, we also trained a statistical model on PCA-diagonalized data using logistic regression model as well as a random forest classifier trained on the untransformed feature-set. When PCA components were used to generate logistic regression models, only PC1 had a significant contribution to the model power, either via single feature permutation test implemented by permutation_importance from scikit-learn, or when it was the sole feature in the model (**Supplemental Figure S8C-E**). The topmost feature of PC1 was cell volume (**Supplemental Figure S7C**), and when we examined how much of total cell volume weight was placed into each PC, PC1 accounted for nearly 40% of cell volume’s total weight (**Supplemental Figure S8F**). When cell cycle classification was performed with a random forest classifier on the original feature-set, the model performed similarly to logistic regression models (**Supplemental Figure S8G**). Similar to the logistic regression model, random forest models where a single-feature was permuted or only a single feature was included, also reveal that cell volume is the most important feature (**Supplemental Figure S8H,I**). Therefore, both the PCA-logistic regression model and the random forest models are consistent with our conclusion that cell volume is likely the major contributor to the accurate classification of cell cycle phase in basal layer stem cells.

### Laser ablation of single cells

The selective ablation of single cells was done similarly to previous methods.^32^ A two-photon laser was tuned to 810nm and used to continuously image a diffraction-limited spot in the middle of a cell nucleus with 20-30% of the total laser power for 15s. For each experiment, 4-6 cells spaced approximately 100µm apart were picked for ablation. Before and after each ablation, an image was taken using normal laser settings to ensure that the intended target received the laser dose. Nuclear volume growth curves were measured as described above, and an exponential growth rate was determined using curve_fit from scipy, assuming an exponential growth function.

### Software

In addition to specific software or libraries noted elsewhere, all computation was done using Python libraries NumPy, SciPy, and Pandas.^67–70^ Visualization was done using matplotlib and seaborn.^71,72^ 3D cell visualization was done using napari.

All code is available at: https://github.com/xies/mouse_skin_size_control

## Supplemental Figure Legends

**Supplemental Figure S1.**
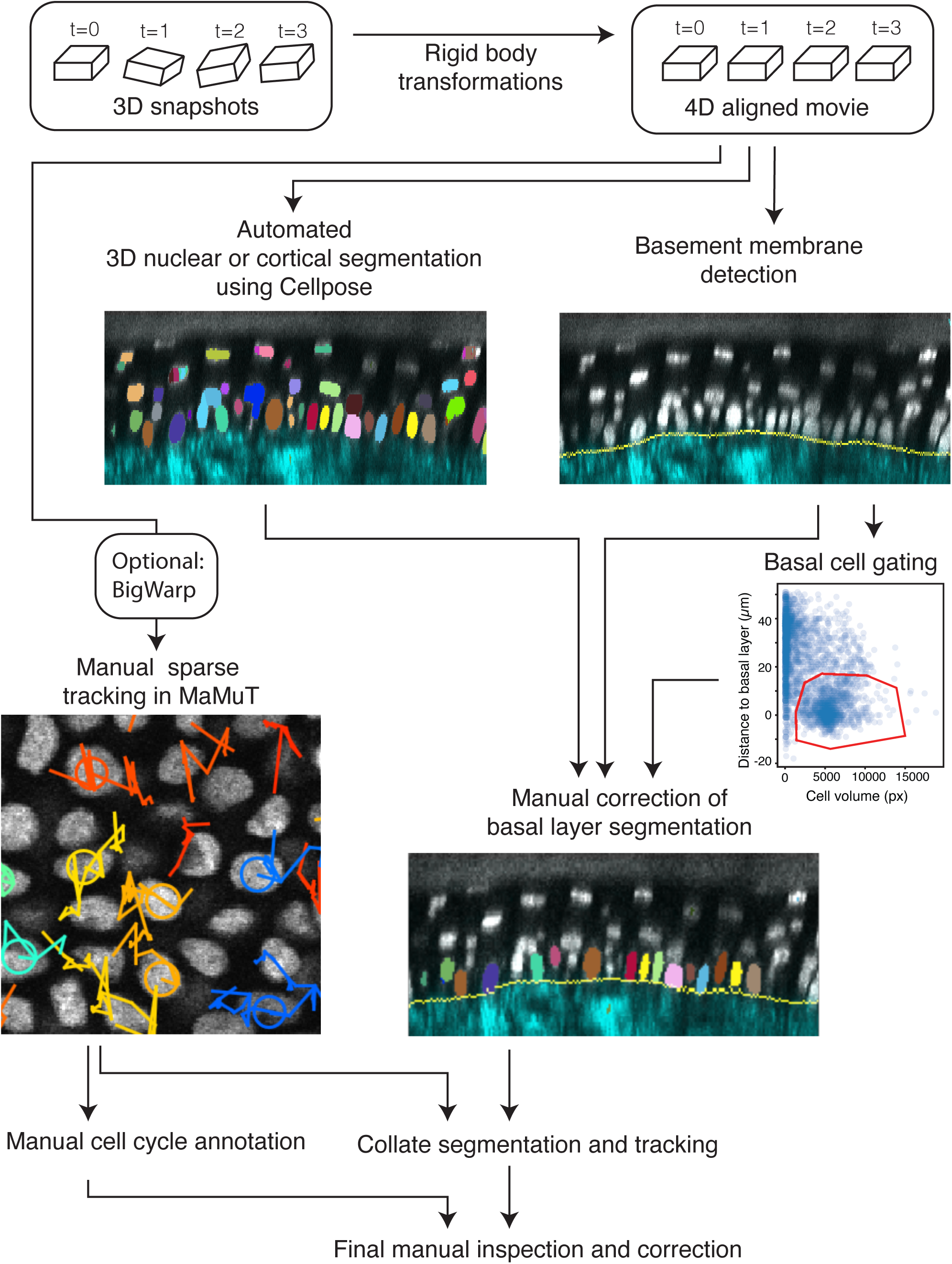
Image analysis pipeline for tracking single cells from longitudinal intravital imaging experiments. A flow chart showing the major steps of the image analysis process used to track single epidermal basal cells through complete cell divisions. Details can be found in Methods.

**Supplemental Figure S2.**
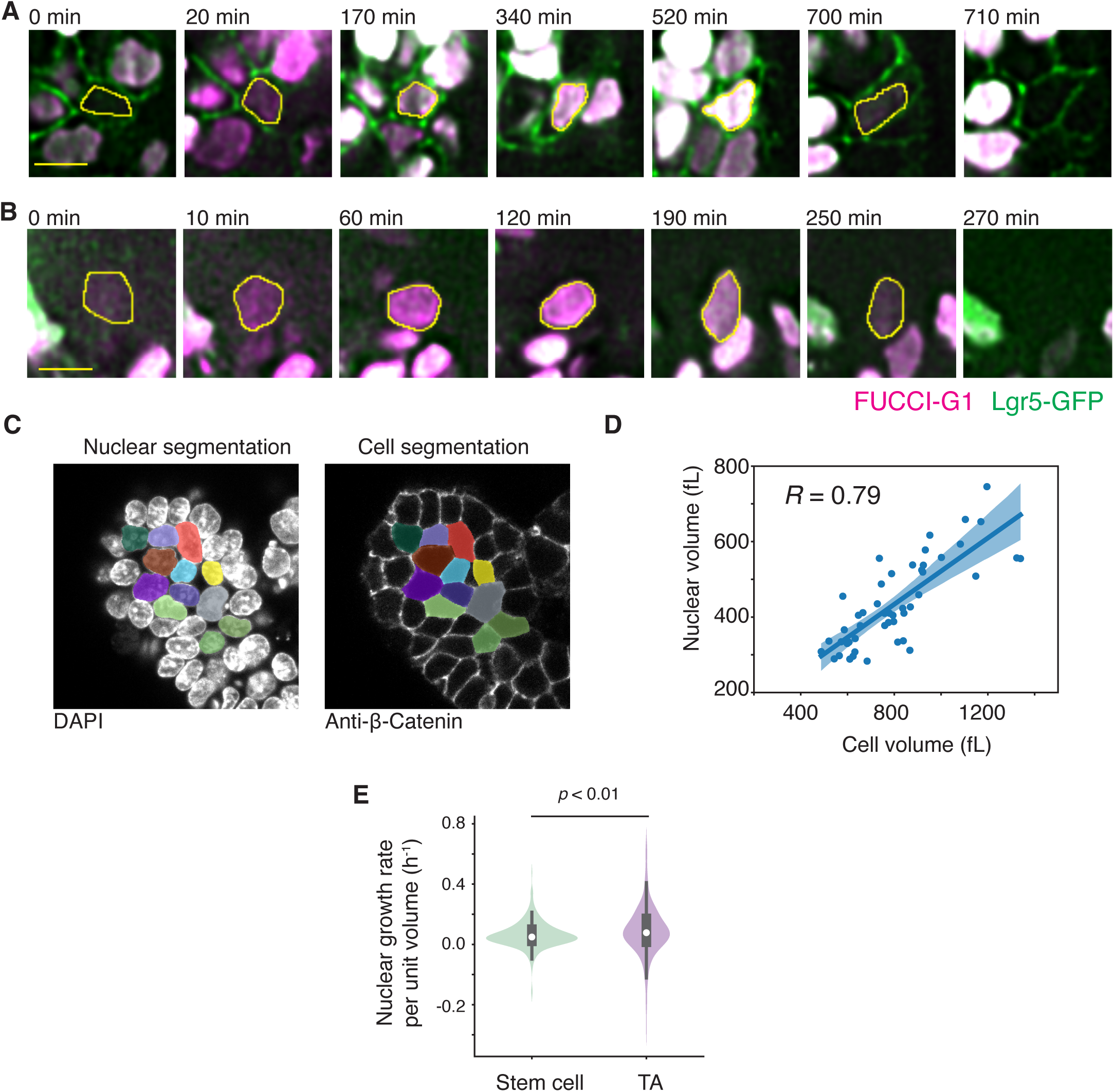
Cell cycle dynamics, nuclear size, cell size, and cell growth in intestinal organoids. A. Example montage of an Lgr5+ intestinal stem cell tracked from birth to G1 exit. Scale bar is 5µm. B. Example montage of an Lgr5-TA cell tracked from birth to G1 exit. C. Example images of nuclear and cell volume segmentation of fixed intestinal organoids based on DAPI and β-Catenin signals, respectively. D. Quantification of the relationship between cell volume and nuclear volume in fixed intestinal organoids. Pearson’s correlation is reported. N = 3 organoids, 48 cells. Solid line is linear regression and shaded area is 95% confidence intervals. E. The size-normalized nuclear growth rate is shown for stem and TA cells. N = 3 organoids, and >250 time points for each cell type. Median is shown in white. Box shows quartile ranges. *p*-value is a two-sample T-test.

**Supplemental Figure S3.**
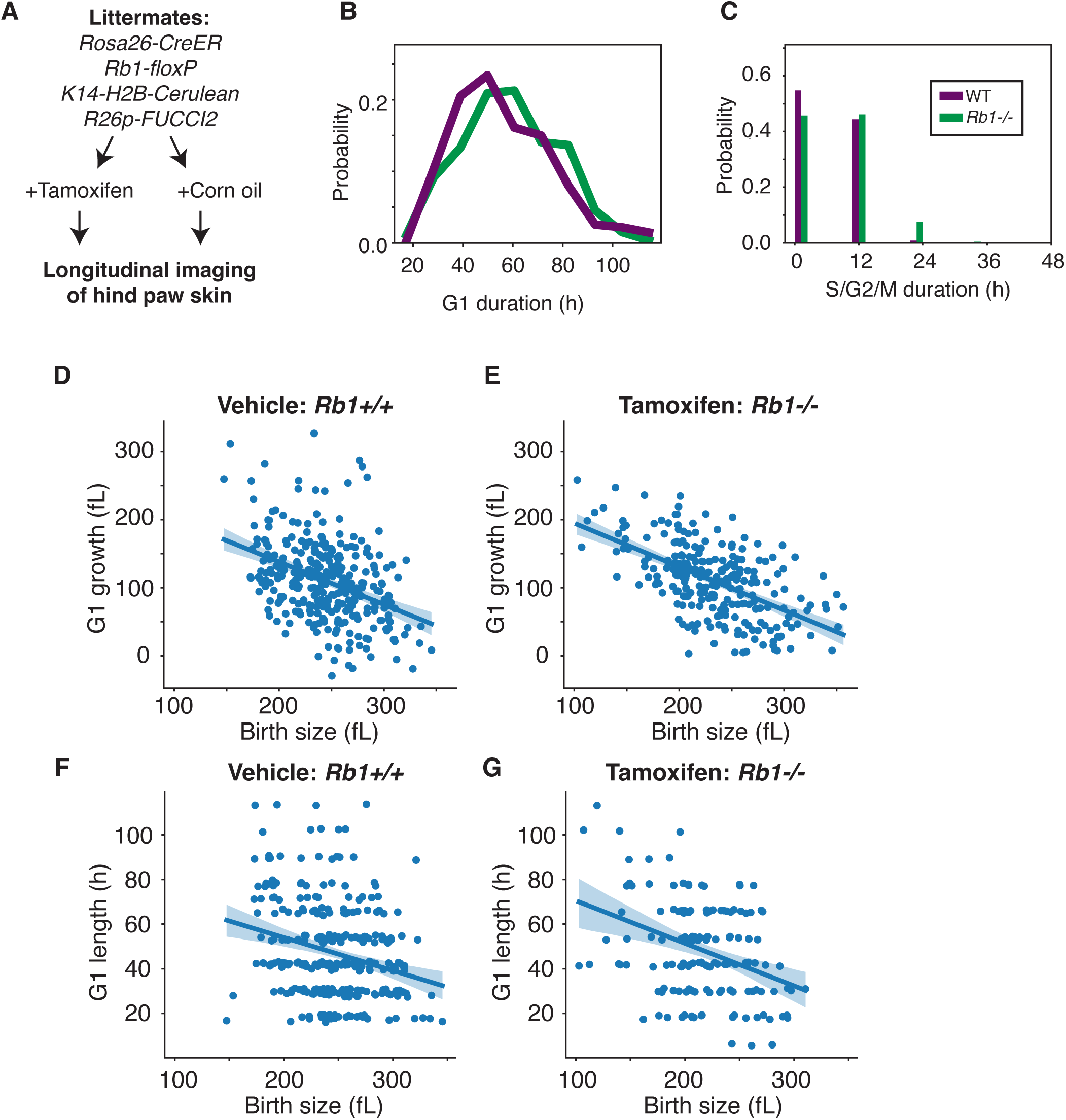
*Rb1−/−* single mutation does not affect cell size control. A. Schematic of the experimental setup. *Rb1-flox; Rosa26-CreERT2; K14-H2B-Cerulean; R26p-FUCCI-G1-mCherry* litter mates were either injected with tamoxifen or corn oil vehicle control, and their paw skin epidermis were longitudinally imaged. B-C. The durations of G1 phase (B) or S/G2/M phases combined (C) are shown for WT and *Rb1−/−* cells. Note that since the sampling rate is every 12h, many cells with S/G2/M phases that were shorter than 12h were annotated to have S/G2/M durations of 0h. D-E. The relationship between nuclear size at birth and amount of nuclear growth during G1 is shown for WT (D) and *Rb1−/−* (E) cells. WT: slope = −0.63; N = 2 mice, 317 cells; *Rb1−/−*: slope = −0.63, N = 2 mice, 244 cells. F-G. The relationship between nuclear size at birth and the duration of G1 phase is shown for WT (F) and *Rb1−/−* (G) cells. WT: Pearson’s R = −0.25; *Rb1−/−*: Pearson’s R = −0.18. For visual clarity, a small random jitter was added to the G1 length. Solid line is linear regression and shaded area is 95% confidence intervals.

**Supplemental Figure S4.**
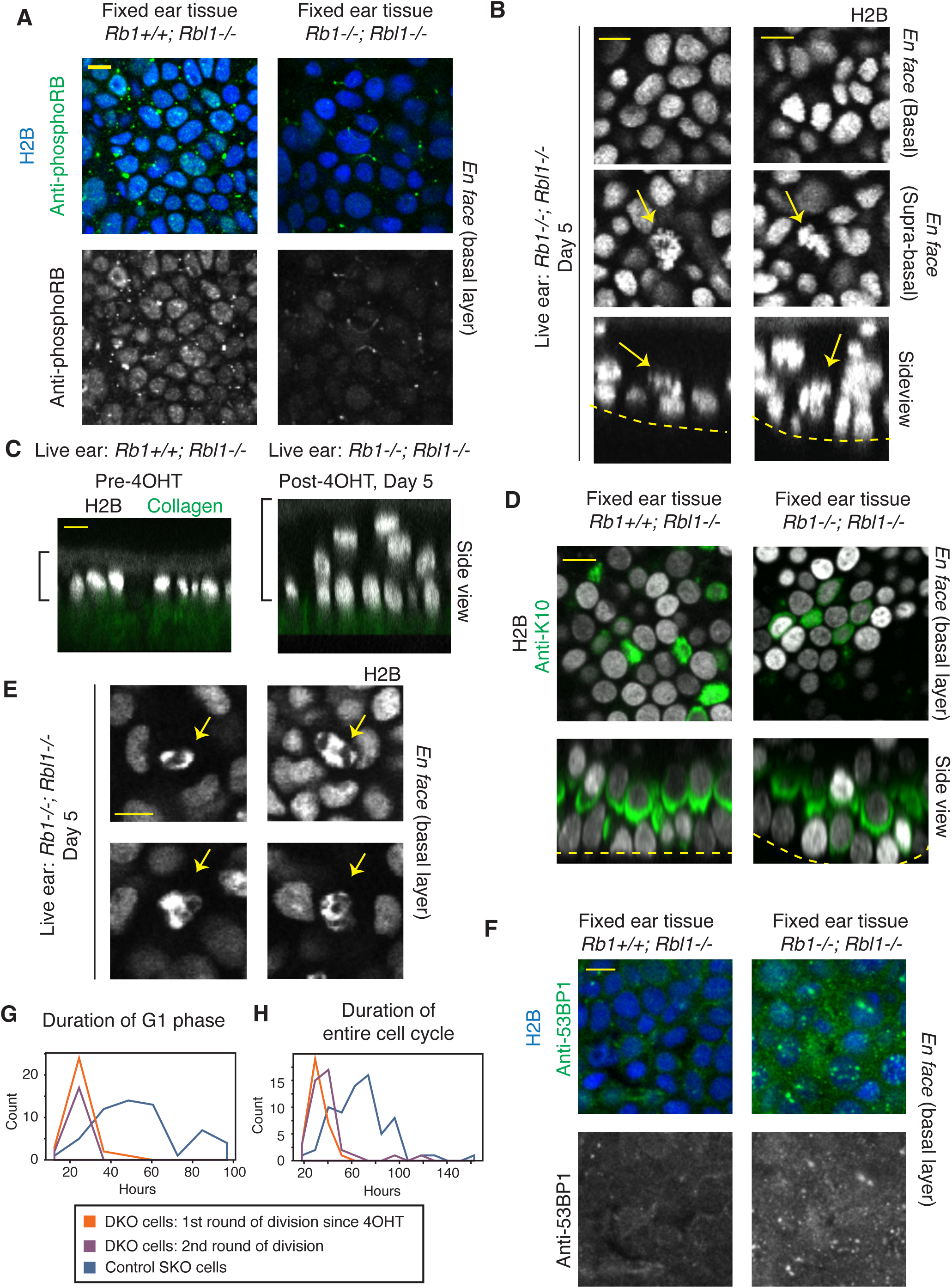
Characterization of *Rb1−/−; Rbl1−/−* DKO tissues. A. Immunofluorescence staining using antibody recognizing phosphorylated Rb1 in fixed *Rb1+/+; Rbl1−/−* (SKO) and *Rb1−/−; Rbl1−/−* (DKO) tissues. An *en face* view of the basal layer is shown. The topical application of 4OHT to the ear skin led to the complete loss of phosphorylated Rb1 signal in DKO tissues. B. Representative images of two regions in the DKO tissue where supra-basal mitotic figures could be seen. The H2B-Cerulean reporter was imaged in a live mouse 5 days after 4OHT treatment. *En face* views of the basal and supra-basal layers and side views are shown. Arrows indicate the location of ectopic mitotic figures. Dotted line indicates the location of the basement membrane. C. Representative side views of an ear epidermis showing the thickening of the epidermis before (left) and after (right) induction of DKO. Bracket denotes the thickness of the epidermis. In the post-4OHT tissue, the indicated epidermal thickness is likely an underestimate. D. Keratin10 (K10) staining in fixed SKO and DKO tissues. Dotted line indicates the location of the basement membrane. E. Representative images of the H2B-Cerulean reporter in DKO live tissues with dead nuclear bodies indicated with arrows. F. Representative images of SKO or DKO fixed tissues stained with 53BP1 antibody. G-H. Histograms of the durations of G1 phase (G) or the entire cell cycle (H) for SKO cells, or the first and second round of division in DKO cells. Scalebar is 10µm.

**Supplemental Figure S5.**
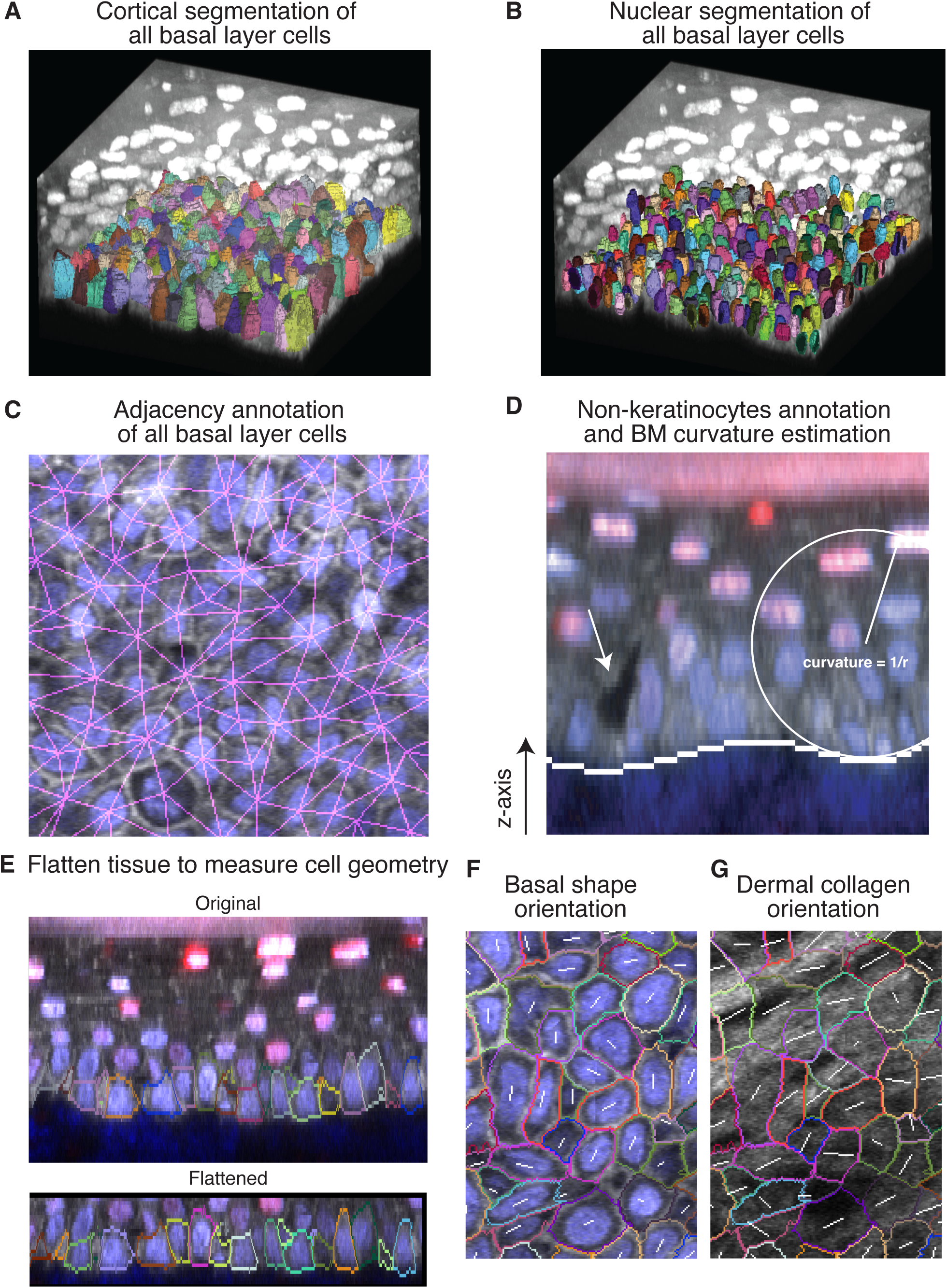
Measurement of cell and microenvironment morphometrics. A. Example image of the exhaustive 3D cell segmentation of all basal layer cells within a ∼120µm x 120µm × 100µm tissue volume. B. Example image of the exhaustive 3D nuclear segmentation of all basal layer cells. C. Example annotation of basal layer cell adjacencies, with each cell-cell interface denoted in magenta. D. Example annotation of basement membrane (BM) location (solid line), calculation of BM curvature (white circle), and annotation of the location of a non-keratinocyte cell shadow in the epidermis (arrow). E. Example of a curved epidermal tissue image being re-sliced to generate a flattened version of the tissue. F. Example measurement of basal shape derived from flattened versions of basal layer cells. Colored outlines show the basal shape segmentation on top of a flattened basal image data. The angle of the white line shows the orientation of the basal shape. The length of the white line shows the eccentricity of the shape. G. Example measurement of the orientation of dermal collagen fibrils. The same basal cell shapes are shown as in (F). The angle of the white line shows the mean orientation of the fibrils within the basal shape. The length of the white line shows the coherence of the fibril orientations within the shape.

**Supplemental Figure S6.**
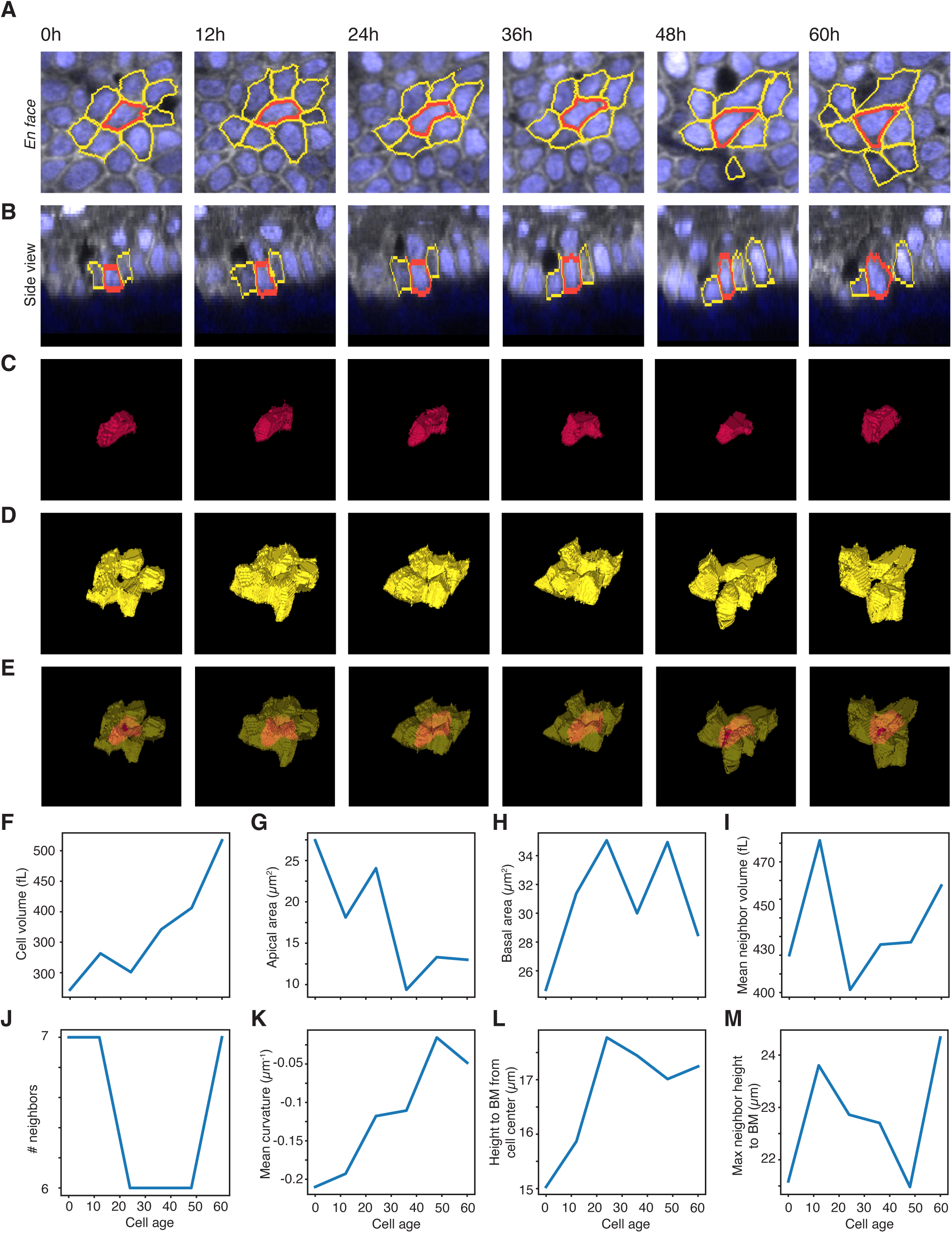
Example cell and microenvironment morphometrics. A-B. The 3D cell segmentations of a central cell (red) and its neighboring cells (yellow) are shown in *en face* (A) and as a side view (B). The cluster of cells are tracked from cell birth to cell division. C-E. A montage of the 3D rendering of the same central cell (C) and its neighboring cells (D) are shown. The merged montage is shown in (E). F-M. For the same central cell, the time-series of its cell volume (F), apical area (G), basal area (H), mean neighbor volume (I), number of neighbors (J), local tissue curvature (K), cell height from basement membrane (L), and maximum neighbor height to basement membrane (M) are shown.

**Supplemental Figure S7.**
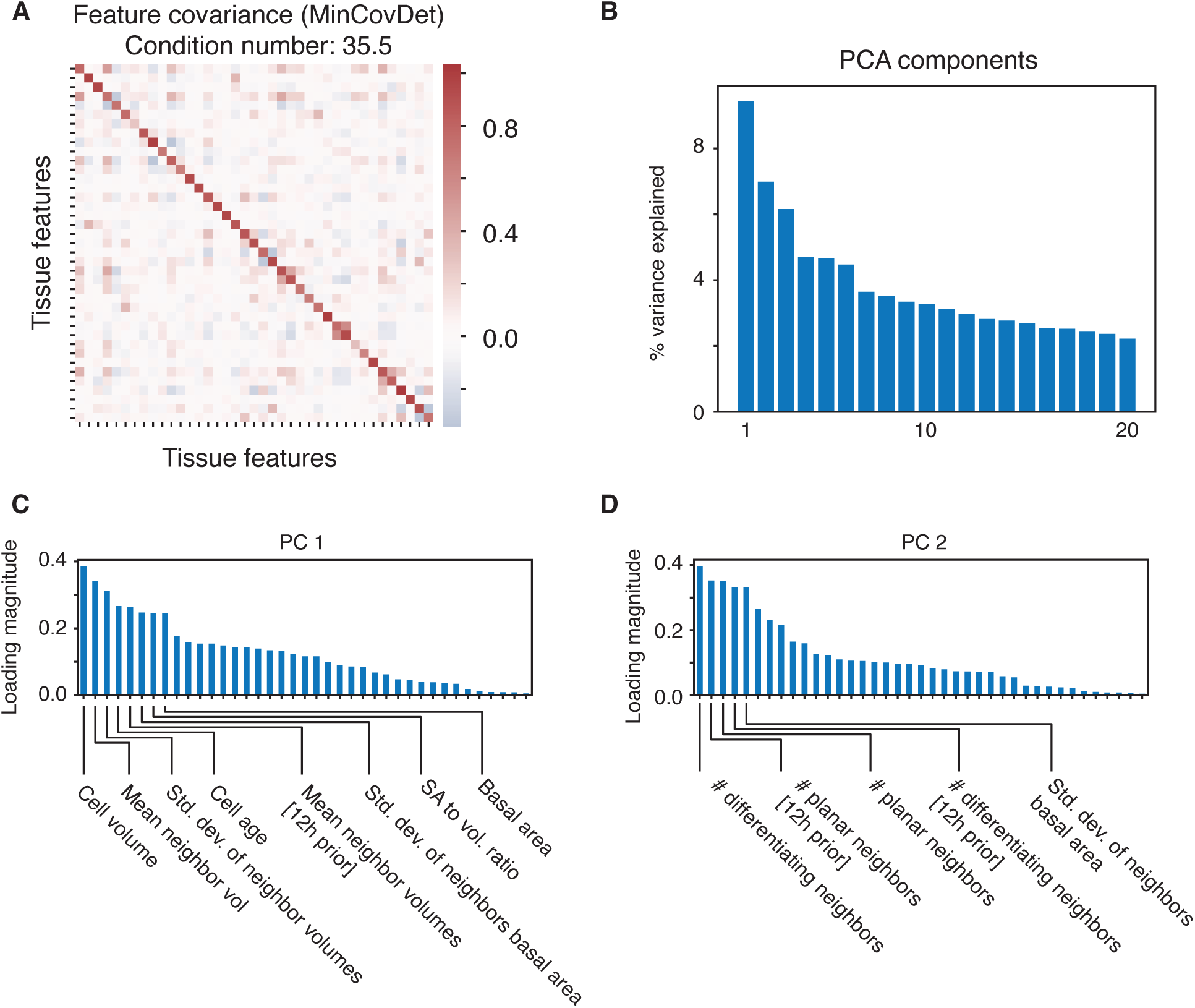
Cell and microenvironment morphometric feature-set. A. The covariance matrix of the feature set used for statistical modeling estimated using the minimum covariance determinant (MinCovDet) is shown (n = 707 samples). The condition number of invertibility, which is the ratio of the largest and smallest eigenvalue, is presented. B. The percent of total variability explained is shown for each component of the data resulting from the principal component decomposition with 20 components. C-D. The top contributing features to the top principal components PC1 (C) and PC2 (D) are shown.

**Supplemental Figure S8.**
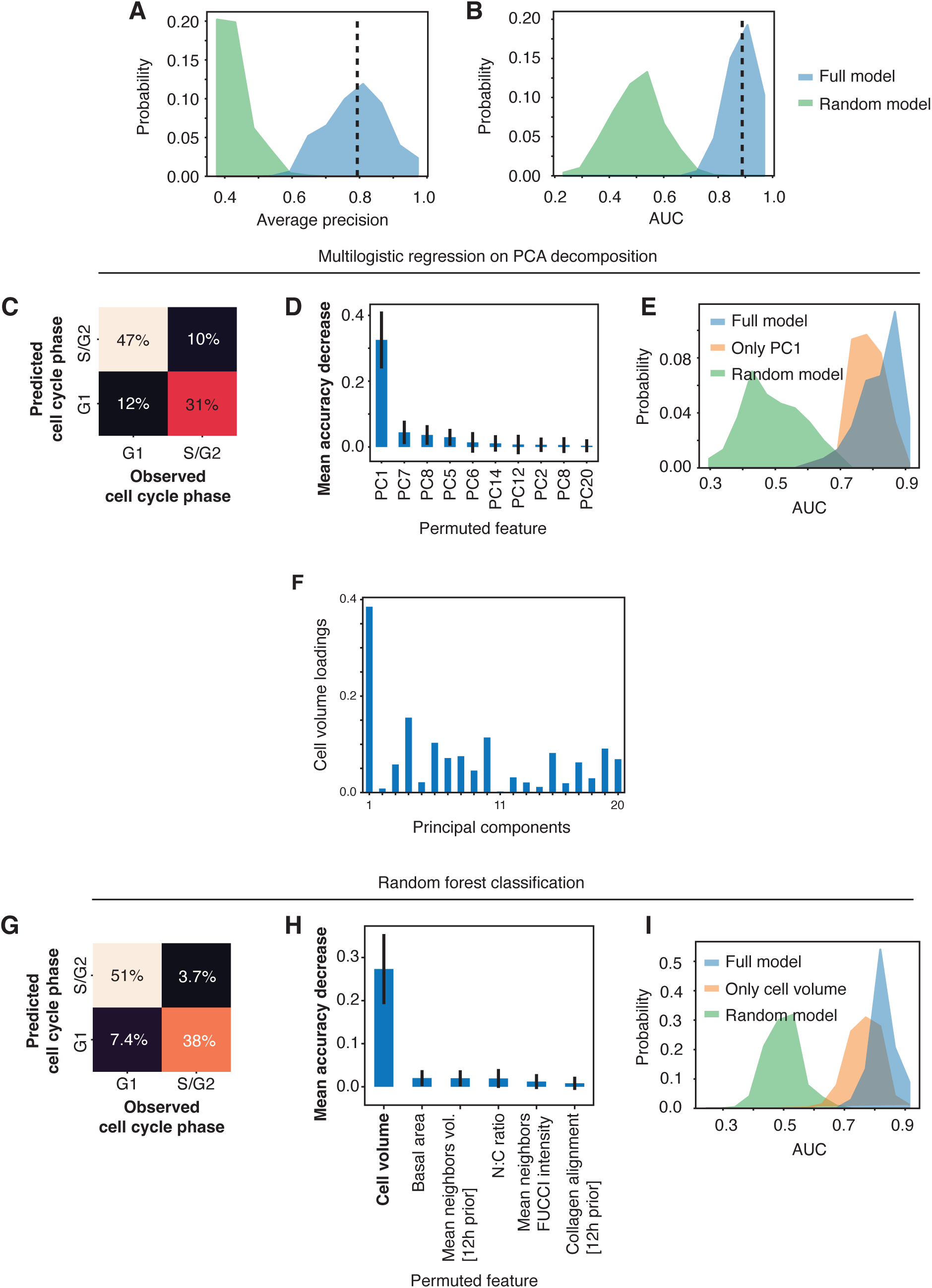
Statistical modeling of cell cycle variability in basal layer epidermal cells. A-B. Histograms of the average precision (A) or AUC (B) of 1000 iterations of cross-validation (10% withheld test data) are shown for the full model or a model of the same size but generated from random data obtained from gaussian distributions. The mean performance of the full model is shown in dotted line. C. The error matrix of a logistic regression model trained on PCA-diagonalized data is shown. D. The relative model accuracy decrease is shown for logistic regression models trained on PCA-diagonalized data, where each indicated principal component was randomly permuted. Error bars indicate std. E. The AUC metric is shown for logistic regression models trained on the full features, random features, or only PC1. F. The loadings of cell volume as a feature onto each PC is shown. G. The error matrix of a random forest model trained on the original data is shown. H. The relative model accuracy decrease is shown for random forest regression models trained on the original data, but where each indicated feature was randomly permuted. Error bars indicate std. I. The AUC metric is shown for logistic regression models trained on the full features, random features, or only on cell volume.

**Supplemental Figure S9.**
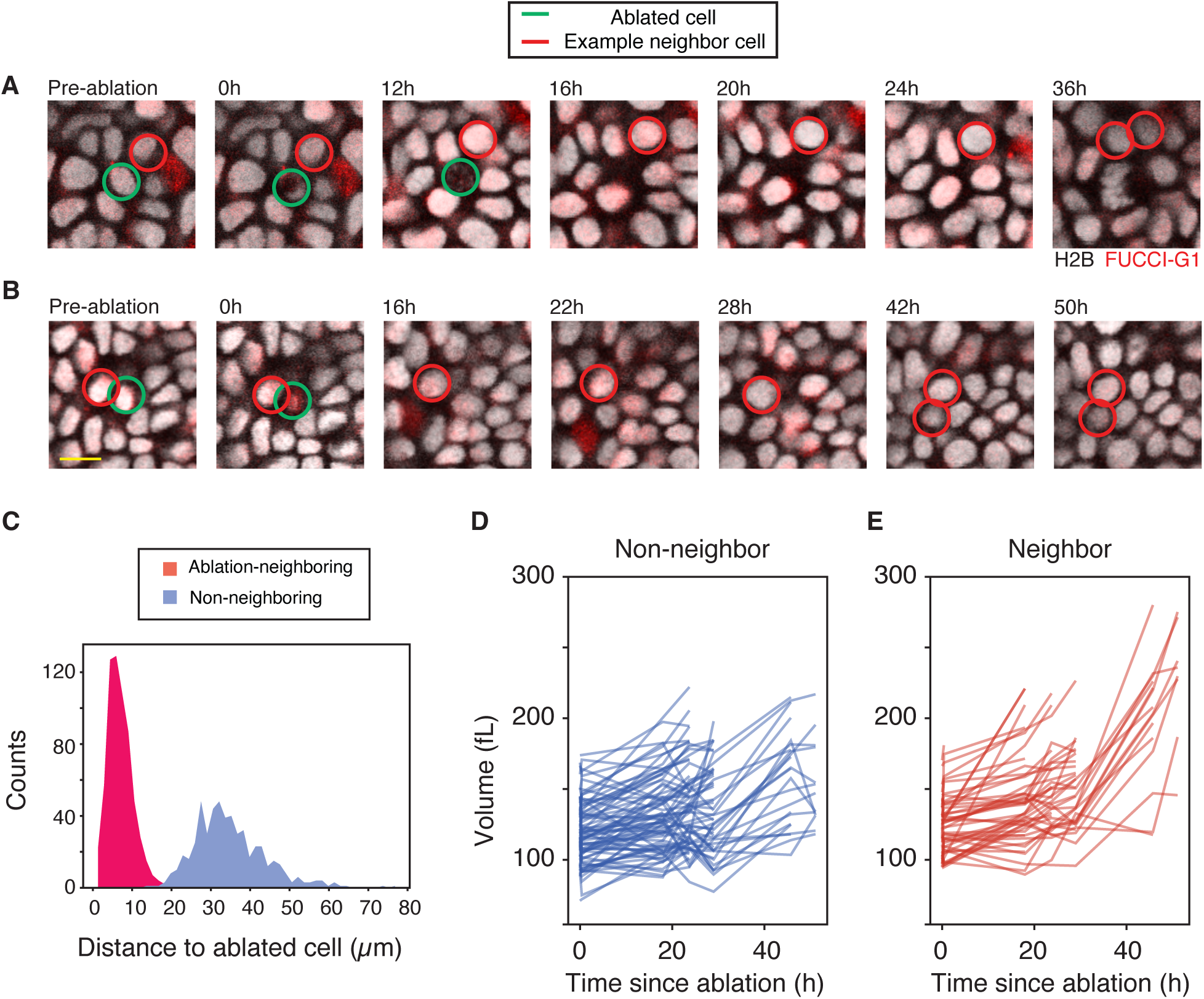
Single cell ablation. A-B. Example montages of cell ablation time-series, showing central ablated cells (green) and single neighbor cells (red) tracked through time. Only a single tracked neighbor cell is highlighted in each montage for clarity. Scale bar is 10µm. Cells were ablated using a two-photon laser as described in the Methods. C. The distributions of distances to the ablation site from the centroids of ablation-neighboring and non-neighboring cells are shown. D-E. The nuclear volume growth curves are shown in a representative ablation experiment, showing cells that neighbored the ablation site (D) and those that were farther away (E). N = 44 non-neighbor, 37 ablation cells

**Supplemental Movie S1-2.** Representative movies of ear epidermis tissues bearing *Rbl1−/−* single mutation (S1) or *Rb1−/−; Rbl1−/−* double mutation (S2) imaged every 12h for 8 days. The movie shows the tissue after the application of ethanol at 0h. A z-slice corresponding to the basal layer is shown at each time point. *K14-H2B-Cerulean* is shown in gray and *Cdt1-mCherry* FUCCI-G1 reporter is shown in red. Single cells are tracked from when they are born until either they exit G1 phase or until they divide into two daughter cells.

**Supplemental Movie S3.** An example 3D segmentation of all basal layer cells within a ∼100µm × 100µm × 100µm in the mouse epidermis. *K14-H2B-Cerulean* is shown in blue and *K14-Actin-GFP* is shown in gray. The cortical and nuclear segmentations for each basal layer are shown in overlay as uniquely colored outlines.

